# Thalamic Interictal Epileptic and Non-Epileptic Events during NREM Sleep in Patients with Focal Epilepsy: a Stereo-EEG Study

**DOI:** 10.64898/2025.12.11.693651

**Authors:** Hongyi Ye, Kassem Jaber, Alyssa Ho, Lingqi Ye, John Thomas, Xiaowei Xu, Cong Chen, Yihe Chen, Guoping Ren, Matthew Moye, Tamir Avigdor, Irena Doležalová, Petr Klimes, Prachi Parikh, Derek Southwell, Zhe Zheng, Junming Zhu, Shuang Wang, Birgit Frauscher

## Abstract

**Background:** Thalamic recordings are increasingly incorporated into stereo-electroencephalography (SEEG) evaluations of drug-resistant focal epilepsy to guide neuromodulation targeting. Human thalamic electrophysiology, however, is poorly defined, limiting the distinction between pathological and physiological activity. Here, we characterized interictal epileptic and non-epileptic events during non-rapid eye movement (NREM) sleep across multiple thalamic nuclei and examined their relationship to seizure outcomes.

**Methods:** We analyzed NREM sleep SEEG recordings from 64 patients with drug-resistant focal epilepsy. Electrodes sampled four thalamic nuclei: centromedian (CM), pulvinar (Pu), ventral lateral (VL), and ventral posterolateral (VPL). Patients were classified into three outcome groups: favorable, unfavorable, and surgically non-remediable. Rates of thalamic spikes, high-frequency oscillations (HFOs), spike-fast activity, and sleep spindles were analyzed and compared across nuclei and outcomes.

**Findings:** Recordings of the thalamus revealed both pathological and physiological interictal events. Interictal epileptic events were infrequent. Only ∼0.2% of seizure-onset zone spikes propagated to the thalamus. Thalamic spike-fast activity was indicative of unfavorable surgical outcomes (CM: *p* = 0.047, *d* = 0.46) or surgically non-remediable epilepsy (VL: *p* = 0.002, *d* = 0.84). In contrast, thalamic sleep spindles were ubiquitous but reduced in surgically non-remediable patients (CM: *p* = 0.031, *d* = –0.58; VL: *p* = 0.005, *d* = –0.79). Finally, unique thalamic SEEG patterns were identified, including spikes concomitant with spindles, isolated spikes, and physiological fast ripples.

**Interpretation:** This study provides a comprehensive characterization of thalamic interictal events during NREM sleep, enriching our understanding of thalamic pathophysiology and highlighting the value of thalamic recordings in presurgical evaluation.

## 1 Introduction

Epilepsy is a network disorder,^1,2^ with focal epilepsy also increasingly understood as a cortico-subcortical network disease.^3^ Both experimental animal and human studies have demonstrated active involvement of the thalamus in epileptic networks, suggesting its function as a key hub for the amplification and propagation of seizures.^4–10^ Clinical trials have shown that thalamic neuromodulation offers a promising treatment for patients with drug-resistant epilepsy who are not candidates for surgically eliminating a defined seizure focus.^11–13^ These findings underscore the growing importance of characterizing electrophysiological properties of the human thalamus and understanding its interactions with cortical networks, as such insights may elucidate the role of specific thalamic nuclei in human epilepsy.

Recently, the use of stereo-electroencephalography (SEEG) to record thalamic activity in patients with epilepsy has become more common.^14,15^ Ictal studies have revealed that the pulvinar nuclei (Pu) may be recruited earlier than the anterior thalamus during seizures.^16,17^ In a cohort of 74 patients, high thalamic epileptogenicity quantified during seizures was associated with poor surgical outcomes.^7^ These findings suggest that multisite thalamic recordings may hold potential not only for guiding neuromodulation targeting and programming but also for providing prognostic information regarding surgical outcomes.

Beyond seizures, interictal EEG events can provide important insights into both pathological and physiological brain activity and may serve as a surrogate marker of epileptogenicity without requiring the recording of seizures. Among them, epileptic spikes are classic interictal epileptiform discharges, while high-frequency oscillations (HFOs) and spike-gamma have emerged as potentially more specific biomarkers of the epileptogenic tissue.^18–21^ These interictal events are optimally analyzed during non-rapid eye movement (NREM) sleep. In this state, artifacts are minimized on SEEG, while spikes and HFOs occur more frequently and propagate more reliably.^22–24^ Moreover, characteristic NREM sleep signatures, such as spindles, provide a window into physiological brain function and help contextualize potential pathological findings.

Despite recent progress, our understanding of thalamic interictal electrophysiology remains limited. While thalamic interictal spikes have been described^25,26^ and their higher rates linked to poor surgical outcomes,^27^ more specific epilepsy biomarkers such as spike-HFOs and spike-gamma have never been systematically examined in the thalamus. The temporal relationship between thalamic spikes and seizure-onset zone (SOZ) discharges also remains largely undefined. Importantly, most prior work has been restricted to a single nucleus ^26–28^, leaving us without a comparative framework across multiple thalamic regions. In addition, only recently has evidence emerged that the thalamus generates distinct physiological SEEG patterns during sleep that differ from cortical activity.^29–31^ Without careful characterization, these physiological patterns risk being misclassified as pathological, underscoring how little is known about the true electrophysiological signatures of the human thalamus.

To address these gaps in knowledge, we investigated interictal SEEG activity in four thalamic nuclei: the centromedian nucleus (CM), Pu, ventral lateral nucleus (VL), and ventral posterolateral nucleus (VPL). With the exception of the Pu, these nuclei have received limited attention in prior human intracranial EEG research. In this study, we specifically focused on the recordings during NREM sleep to leverage the optimal signal quality described above. Our primary objective was to establish benchmarks for interictal thalamic SEEG interpretation by characterizing the electrophysiological profiles and rates of interictal epileptic and non-epileptic events across multiple thalamic nuclei, as well as assessing the co-occurrence of spikes between the thalamus and the SOZ. Secondarily, we aimed to investigate the relationship between these thalamic features and epilepsy prognosis. We hypothesized that the extension of the interictal epileptic network into the thalamus, manifested by higher burdens of epileptiform discharges, would be associated with poor seizure outcomes.

## 2 Methods

### 2.1 Patient Selection

This study included patients with drug-resistant focal epilepsy who underwent SEEG monitoring for epileptogenic zone localization with at least one electrode inserted in the thalamus at the Second Affiliated Hospital of Zhejiang University (SAHZU) between March 2022 and December 2023 and at the Duke University Medical Center (DUMC) between March 2023 and June 2024. All patients underwent comprehensive presurgical evaluations, including detailed clinical history, neuroimaging, neuropsychological assessments, and video-scalp EEG monitoring. Individualized electrode implantation plans were formulated during the multidisciplinary presurgical epilepsy conference, based on findings from these evaluations. Patients receiving subsequent resective or ablative surgery underwent regular follow-up exceeding one year, with surgical outcomes classified using the Engel scale (Table S1).^32^ Patients deemed ineligible for surgery due to widespread or multi-focal SOZs were marked as “surgically non-remediable”. Exclusion criteria were: (1) all thalamic SEEG channels showing significant signal artifacts; (2) history of previous resective neurosurgery; (3) failure to obtain qualifying interictal NREM sleep recordings due to frequent seizures (see Section 2.3, Fig. S1).

#### Ethics

The study adhered to the principles of the Declaration of Helsinki and the protocol was approved by the institutional review boards of both epilepsy centers (SAHZU: 2024-0750; DUMC: Pro00115576). Informed consent was obtained from all patients or their legal guardians.

### 2.2 SEEG evaluation and contacts localization

Both epilepsy centers used SEEG electrodes with identical specifications (contact diameter: 0.8 mm; contact length: 2 mm; intercontact spacing: 1.5 mm). Sampling rates were 2000 Hz at SAHZU and 2048 Hz at DUMC, with DUMC data downsampled to 2000 Hz for consistency. The SOZs were annotated jointly by epileptologists and clinical neurophysiologists. An example of the electrode implantation scheme is shown in **Fig. 1a**. In our cohort, the CM and Pu were the two primary implanted nuclei during clinical SEEG exploration. Due to anatomical reasons, electrode trajectories often passed through lateral thalamic nuclei, including the VL and the VPL (**Fig. 1b**). Entry points for thalamic electrodes typically included the precentral gyrus, middle frontal gyrus, inferior frontal gyrus, or the superior temporal gyrus. High-resolution contrast-enhanced T1-weighted MRIs were used for preoperative planning, and electrode trajectories were adjusted by neurosurgeons to avoid intracranial vasculature.

**Figure 1.**
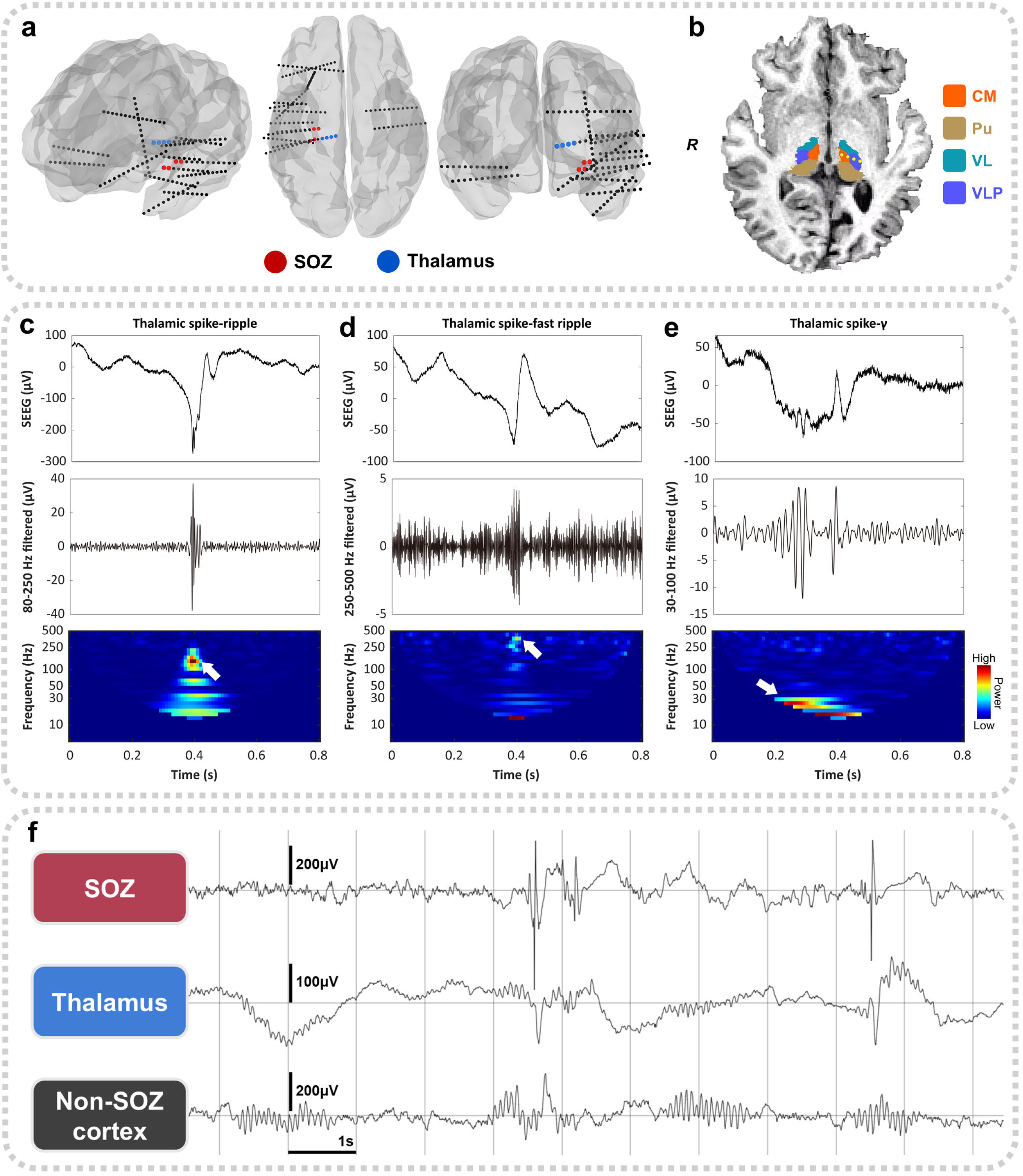
Thalamic electrode implantation and interictal transient events. (a) SEEG electrode reconstruction in a patient, which demonstrates the SEEG implantation scheme. Dots indicate electrode contacts. (b) Relative positions of the four thalamic nuclei studied in our work and SEEG contacts in the MR image of an example patient. The contact markers are schematic representations indicating the location of the sampled nuclei and they were projected onto the segmentation map. (c)-(e) Examples of thalamic spike-ripple, spike-fast ripple, and spike-gamma events, as well as their respective time-frequency spectra. (f) An example of interictal SEEG recordings during NREM sleep. We observed that spikes and spindles can co-occur in the thalamus. SOZ, seizure onset zone; CM, centromedian nucleus; Pu, pulvinar nuclei; VL, ventral lateral nucleus; VPL, ventral posterolateral nucleus; SEEG, stereo-electroencephalography; NREM, non-rapid eye movement.

To localize intracranial electrode contacts, presurgical MRIs and post-implantation CT scans were co-registered and fused. Neuroanatomical segmentations were performed using FreeSurfer.^33^ Each thalamic contact was assigned to a specific anatomical nucleus based on a standardized atlas,^34^ with localization visually confirmed by two reviewers (K.J. and L.Y.).Table S1, S2 summarize the thalamic nuclei sampled across the study population. The CM, Pu, VL, and VPL were the most commonly sampled nuclei and were selected for further analysis (**Fig. 1b**). The CM exhibits widespread connectivity to the cortex, while the Pu maintains extensive connections with occipito-parietal and temporal lobes.^35,36^ In contrast, the VL and VPL function as specialized relays, projecting primarily to the motor and sensory cortices, respectively.^36,37^

The default reference for SEEG signals was the skull electrode at DUMC and was the average signal of two white matter channels at SAHZU. To emphasize local electrophysiological activity and reduce noise and volume effects, a bipolar montage was used for all SEEG analyses. The bipolar channel coordinate was defined as the midpoint between the two contacts. If one contact was fully within the nucleus and the paired contact was located at the boundary, the bipolar channel was still retained.^38^ When multiple bipolar channels sampled the same thalamic nucleus within a patient, the one closest to the center of the nucleus was selected for analysis.

### 2.3 NREM sleep recordings

Previous studies have shown that NREM sleep during the first sleep cycle generally contains the highest density of interictal epileptic events and exhibits fewer artifacts compared to wakefulness.^39,40^ Thus, NREM sleep SEEG recordings from the first sleep cycle were selected for analysis. In 54 patients, scalp and intracranial EEG were recorded simultaneously. Subdermal needle electrodes were placed at F3, F4, Fz, C3, C4, Cz, T3, T4, P3, and P4. Additional disc electrodes were placed on bilateral mastoids, and we also recorded electrooculogram (EOG) and electromyogram (EMG) signals. Scalp EEG, EOG, and EMG were referenced to the mastoids, and 30-second epochs were manually scored according to the American Academy of Sleep Medicine criteria.^41^ In this study, NREM sleep refers to the combination of N2 and N3 stages.^42^ For ten patients without simultaneous scalp EEG recordings, sleep stages were automatically classified using our openly available SleepSEEG toolbox,^24^ which has demonstrated reliable performance in identifying NREM sleep based on intracranial EEG.

For each case, a 20–60-minute segment of continuous NREM sleep was selected from the first sleep cycle, based on the following criteria: (1) at least 48 hours post-implantation; (2) at least 4 hours before and after the most recent clinical seizure; (3) at least 15 minutes before and after the most recent subclinical pure EEG seizure. If no qualifying segment was available, the patient was excluded from the study. If multiple qualifying segments existed in a patient, the longest one was selected. This duration range was chosen to maximize the capture of interictal events while accommodating variability of duration of the first sleep cycle.

### 2.4 Interictal event detection

During NREM sleep, typical interictal transient events recorded via SEEG include spikes, HFOs, and sleep spindles. HFOs are categorized as ripples (80–250 Hz) and fast ripples (250–500 Hz), and may represent either physiological or pathological activity when recorded in the thalamus.^29,43^ Although spikes are usually considered interictal epileptiform discharges in patients with epilepsy, their identification is subjective and exhibits low inter-rater reliability.^44^ To improve specificity for epileptic events, we incorporated two additional biomarkers: spike-HFO and spike-gamma. Spike-HFO refers to the co-occurrence of a spike and an HFO, either in the form of a spike-ripple (**Fig. 1c**) or a spike-fast ripple (**Fig. 1d**), which have been validated as reliable epileptiform markers.^20,45,46^ Spike-gamma is defined as a spike preceded by at least three cycles of gamma-band oscillations (30–100 Hz) within 200 ms before spike onset (**Fig. 1e**).^47^ Recent studies have shown that spike-gamma events outperform isolated spikes in localizing the epileptogenic zone.^18,21^ In the present study, spike-HFO and spike-gamma events were grouped into a single category, termed spike-fast activity. Sleep spindles (10–16 Hz) are characteristic physiological features of NREM sleep (**Fig. 1f**).

Interictal SEEG transient events were detected using automated algorithms implemented in our established analysis pipeline, which has been utilized in sleep and epilepsy research.^21,42,48^ Spikes were detected using an adaptive algorithm that models the statistical distribution of signal envelopes to distinguish discharges from background activity.^49^ The gamma component within spike-gamma events was characterized using time-frequency analysis via continuous wavelet transforms.^21,47^ Spindles were identified based on root-mean-square amplitude thresholds of band-pass filtered signals.^42,50^ For HFOs, a multi-narrowband filter bank approach was utilized to capture localized power increments relative to a moving background baseline.^51^ The corresponding open-source code is listed in Table S3. Spike-HFO events were labeled by identifying overlaps between detected spikes and HFOs.

Since detection parameters for all interictal event detectors were originally set for cortical recordings, we first evaluated their performance in thalamic data. Five patients with temporal, frontal, posterior cortex, insular, or multi-lobar epilepsy were randomly selected. For each, five minutes of thalamic SEEG data during NREM sleep were manually annotated for spikes, HFOs, and spindles. Detector outputs were then compared against manual labels. While spindle and HFO detectors showed comparable sensitivity and precision to their cortical performance, spike detectors exhibited reduced precision, often misclassifying spindles as spikes (Figs. S2, S3). Therefore, all automatically detected thalamic spikes were manually reviewed to exclude mislabeled spindles and those that did not stand out from the background. Spindle markers overlapping with spikes were then visually verified, and among these markers only those in which spindles and spikes did co-occur (**Fig. 1f**) were retained. To assess the reliability of this approach, we conducted an additional two-round validation against a human expert (gold standard) in a subset of ten patients. This analysis confirmed that while our semi-automated pipeline yielded a more conservative spike count than the raw detector, it was highly selective (median = 1.0), effectively minimizing the inclusion of sleep spindles (see Supplementary Materials and Fig. S3).

### 2.5 Spike co-occurrence between the SOZ and thalamus

We observed that spikes in the thalamus sometimes, but not always, co-occurred with spikes in the SOZ (**Figs. 1f, 2a-c**). A 150-ms temporal window was used to define spike co-occurrence.^52,53^ However, in patients with frequent interictal spikes, co-occurrence may arise by chance rather than reflecting true spike propagation or co-activation. To reduce the impact of spurious co-occurrence, we employed three strategies: (1) For spikes detected across multiple SOZ channels, only the earliest spike marker within a 150-ms co-occurrence window was retained, which reduced the density of spike markers. The same procedure was applied to thalamic spikes across different nuclei. (2) Only patients with a focal SOZ - defined as in our previous work - were included in the co-spike analysis,^54^ as they typically had well-defined SOZ and exhibited fewer SOZ channels as well as spike markers. (3) Surrogate data analysis was used to estimate and correct for chance co-occurrences. Specifically, let n_soz_ and n_th_ represent the number of spikes in the SOZ and thalamus, and let D denote the duration of NREM recordings. Random spike timestamps were generated under a continuous uniform distribution assumption, with n_soz_ and n_th_ events randomly distributed within D to form one surrogate dataset (**Fig. 2d**). The number of co-spikes was computed for each set. This procedure was repeated 5000 times, yielding a distribution of random co-occurrence counts (**Fig. 2d**).

**Figure 2.**
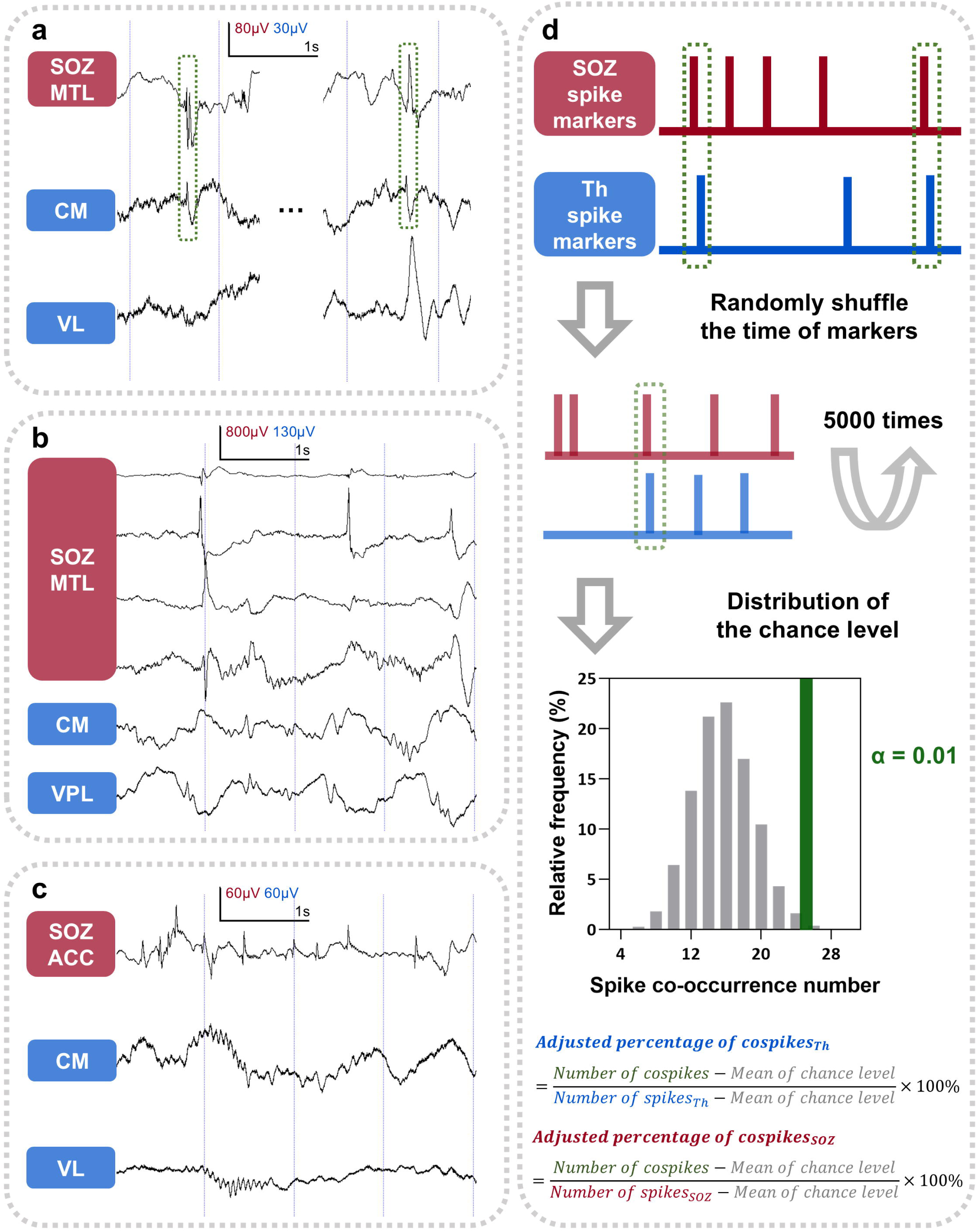
Analysis of spike co-occurrence between the SOZ and the thalamus. (a) Co-spikes between the SOZ and the thalamus. These thalamic spikes are considered as interictal epileptiform discharges. (b) There are spikes within the thalamus, but they do not co-occur with the SOZ ones. (c) No spikes occur in the thalamus while the SOZ is spiking. (d) Spike co-occurrence percentages between the SOZ and the thalamus were corrected by surrogate data analysis to reduce the effect of random co-occurrence. Two co-occurrence percentages can be obtained by using the respective numbers of spikes in the thalamus and SOZ as the denominators. SOZ, seizure onset zone; CM, centromedian nucleus; VL, ventral lateral nucleus; VPL, ventral posterolateral nucleus; MTL, mesial temporal lobe; ACC, anterior cingulate cortex.

We then defined the α-value as the proportion of surrogate datasets in which the number of co-occurrences exceeded or equaled the observed value. A lower α-value indicates that the actual co-spike count is unlikely to result from chance. In addition, the average number of random co-spikes from the surrogate distributions was also calculated. Then, two adjusted percentages of co-spikes were derived using the spike numbers of the SOZ and thalamus as denominators (**Fig. 2d**).

To evaluate directionality of spike propagation between the SOZ and thalamus, a sign test was performed based on the interval of spike co-occurrences.^55^ A *p*-value < 0.05 was considered indicative of significant unidirectional propagation.

### 2.6 Functional connectivity between the SOZ and thalamus

To further investigate the influence of SOZ-thalamus connectivity on the proportion of co-spikes, we analyzed functional connectivity between the SOZ and thalamic nuclei in patients with focal SOZs. We estimated directed connectivity *̑X*_i,j_ using time-domain Granger causality. Preprocessing included bandpass filtering (0.3–512 Hz) with a powerline notch filter on SEEG, followed by detrending and channel-wise scaling. Granger causality was then computed using a 2-minute window with zero overlap for SEEG channel pairs between the SOZ and the thalamus. Let *̑X* denote the patient-specific adjacency matrix, where s represents the set of SOZ channels and *T* represents the set of thalamic channels. An entry *̑X*_i,j_ represents the Granger causal influence from channel *i* (where *i* ɛ *s*) to channel *j* (where i ɛ *T*). This influence quantifies the degree to which past values of channel *i* improve the prediction of channel *j* compared to a univariate autoregressive model of channel *j* alone. The model order was optimized for each signal pair using the Brainstorm implementation.

To quantify the directionality of the interaction between the SOZ and the thalamus, we computed the *asymmetry index* (ASI) as follows:

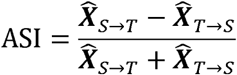

Where 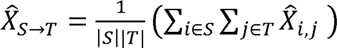 and 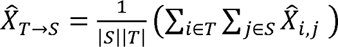 represent the mean directed connectivity from the SOZ to thalamus, and from the thalamus to SOZ, respectively. The ASI ranges from -1 to 1, where -1 indicates dominant connectivity from the thalamus to the SOZ, 1 indicates dominant connectivity from the SOZ to the thalamus, and 0 indicates either symmetric or negligible directed connectivity. Finally, we assessed the relationship between the ASI value and the proportion of co-occurring spikes using Spearman’s correlation coefficient.

### 2.7 Statistical analysis

Normality of all datasets was tested using the Shapiro–Wilk test. As most data were not normally distributed, non-parametric statistical tests were used throughout. Data were reported as median value with interquartile range (IQR). Effect sizes for non-normally distributed variables were computed using *Cliff ’s delta*. For each patient, the median event rate across all sampled thalamic nuclei was calculated to represent the event rate for the whole thalamus. Group comparisons based on seizure outcomes were conducted using the Kruskal–Wallis test, followed by post-hoc pairwise comparisons with Bonferroni correction for the number of groups. Comparisons across thalamic nuclei were performed using paired Wilcoxon signed-rank tests. To compare the adjusted percentage of co-spikes between patients with favorable (Engel I) vs. unfavorable surgical outcomes (Engel II-IV), the Mann–Whitney U test was employed. All statistical analyses were performed using R (version 4.3.3).

#### Role of funders

The funders played no role in the study design, data collection, analysis, interpretation, or the writing of the manuscript. None of the authors have been paid to write this article by a pharmaceutical company or other agency.

## 3 Results

### 3.1 Patient characteristics

The final study cohort consisted of 64 patients (32 females) with drug-resistant focal epilepsy (Fig. S1). Demographic and clinical data are summarized in Table S1. Among the thalamic nuclei sampled, the CM was recorded in 50 (78.1%) patients, the VL in 36 (56.3%), the VPL in 31 (48.4%), and the Pu in 11 (17.2%). Of the 45 patients (70.3%) who underwent epilepsy surgery, 22 achieved favorable surgical outcomes (Engel class I, free of disabling seizures). Sixteen patients (25.0%) were deemed not surgically remediable following SEEG evaluation. Neither resection nor laser ablation was recommended for these patients due to widely distributed or multi-focal SOZs (*n* = 15) or localization of the SOZ within eloquent cortex (*n* = 1). The latter case was excluded from comparisons between outcome groups. An additional three patients (4.7%) were awaiting epilepsy surgery and were not assigned to any outcome group, so they were not included in the outcome comparisons below.

### 3.2 Rates of interictal events vary among outcome groups in the thalamus as a whole and across investigated subnuclei

Across all patients, median rates of thalamic interictal events during NREM sleep were 0.58 (IQR = 0.26–1.32) events per minute for spikes, 0.28 (IQR = 0.15–0.68) for ripples, 0.41 (IQR = 0.11–1.04) for fast ripples, and 3.67 (IQR = 3.16–4.79) for spindles. These rates did not differ significantly among the favorable, unfavorable, and non-remediable groups except for spikes (surgically non-remediable vs. Engel I, *p* = 0.049, *d* = 0.47; **Fig. 3a**). However, spike-fast activity (median rate: 0.07 events/min, IQR = 0.01–0.20), considered a more specific marker of epileptogenicity, exhibited more significant differences across groups. Patients with unfavorable surgical outcomes and those who were surgically non-remediable showed significantly higher spike-fast activity rates in the thalamus compared to those with favorable outcomes (*p* = 0.034, *d* = 0.44; *p* = 0.004, *d* = 0.62, respectively; **Fig. 3a**). To validate the robustness of this association, we performed a multivariate logistic regression adjusting for baseline characteristics, including age, biological sex, and epilepsy duration. The predictive performance was further evaluated using leave-one-out cross-validation. The results indicated that the spike-fast activity rate remained significantly associated with seizure outcomes independent of these demographic factors (Table S4, Fig S4).

**Figure 3.**
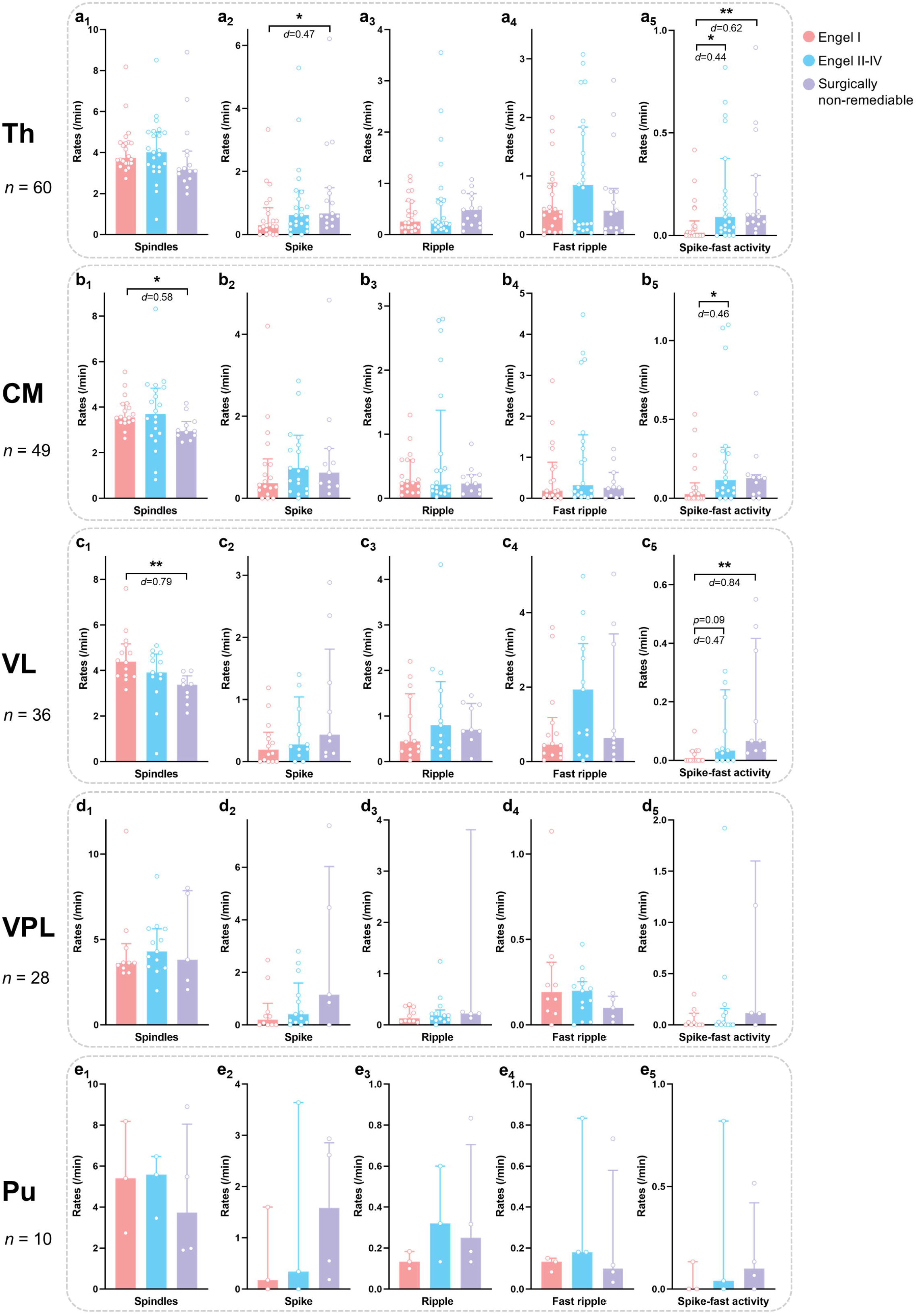
Rates of epileptic and non-epileptic interictal events during NREM sleep in different thalamic nuclei and in three outcome groups. (a) Event rates in the thalamus as a whole. Surgically non-remediable patients have more spike-fast activity and spikes than patients with favorable surgical outcomes. Patients with unfavorable surgical outcomes have more spike-fast activity than those with favorable surgical outcomes. (b) Event rates in the CM. Patients with unfavorable surgical outcomes have more spike-fast activity than patients with favorable surgical outcomes. Surgically non-remediable patients have fewer spindles than those with favorable surgical outcomes. (c) Event rates in the VL. Surgically non-remediable patients have more spike-fast activity but fewer spindles than patients with favorable surgical outcomes. (d)-(e) There was no difference among the three groups in the VPL and the Pu. Median values and interquartile ranges are shown in the plots. Each dot represents one subject. For better visualization, the plots a_2_, a_5_, b_2_, d_3_, and d_5_ each exclude one subject with very high event rates. *, p < 0.05; **, p < 0.01; Th, thalamus; CM, centromedian nucleus; Pu, pulvinar nuclei; VL, ventral lateral nucleus; VPL, ventral posterolateral nucleus.

Nucleus-specific analyses revealed further differences. In the CM, the rate of spike-fast activity was significantly elevated in patients with unfavorable surgical outcomes compared to those with favorable outcomes (*p* = 0.047, *d* = 0.46; **Fig. 3b**). The surgically non-remediable group also showed reduced spindle density in CM compared to the favorable outcome group (*p* = 0.031, *d* = –0.58).

In the VL, patients classified as surgically non-remediable had both significantly higher rates of spike-fast activity (*p* = 0.002, *d* = 0.84) and significantly lower spindle rates (*p* = 0.005, *d* = –0.79) compared to the favorable outcome group (**Fig. 3c**). Although the spike-fast activity rate in the unfavorable surgical outcome group did not reach statistical significance versus the favorable surgical outcome group (*p* = 0.091, *d* = 0.47), the spike-fast ripple rate was significantly elevated (*p* = 0.040, *d* = 0.38; Fig. S5).

For other interictal event types, including spikes, ripples, and fast ripples, no significant group differences were observed in either CM or VL. Moreover, in the Pu and VPL, none of the interictal transient events showed significant differences across outcome groups (Figs. 3d-e).

In a supplementary analysis, no significant correlation was found between the latency of thalamic seizure involvement and the rates of interictal events (Figs. S6a-e).

### 3.3 The density of interictal events varies across thalamic nuclei

To investigate regional differences in thalamic SEEG activity, interictal event rates during NREM sleep were compared across nuclei within individual patients. We found that the VL exhibited significantly higher rates of spindles (*p* = 0.018, *d* = 0.25), ripples (*p* < 0.001, *d* = 0.46), and fast ripples (*p* = 0.007, *d* = 0.37) than the CM (**Fig. 4a**). The CM, in turn, had significantly more fast ripples (*p* = 0.012, *d* = 0.24) and spike-fast ripples (*p* = 0.037, *d* = 0.36) compared to the VPL (**Fig. 4b**, Fig. S7). Due to small sample sizes, statistical comparisons for other nucleus pairs were not performed (Table S1, Fig. S7).

**Figure 4.**
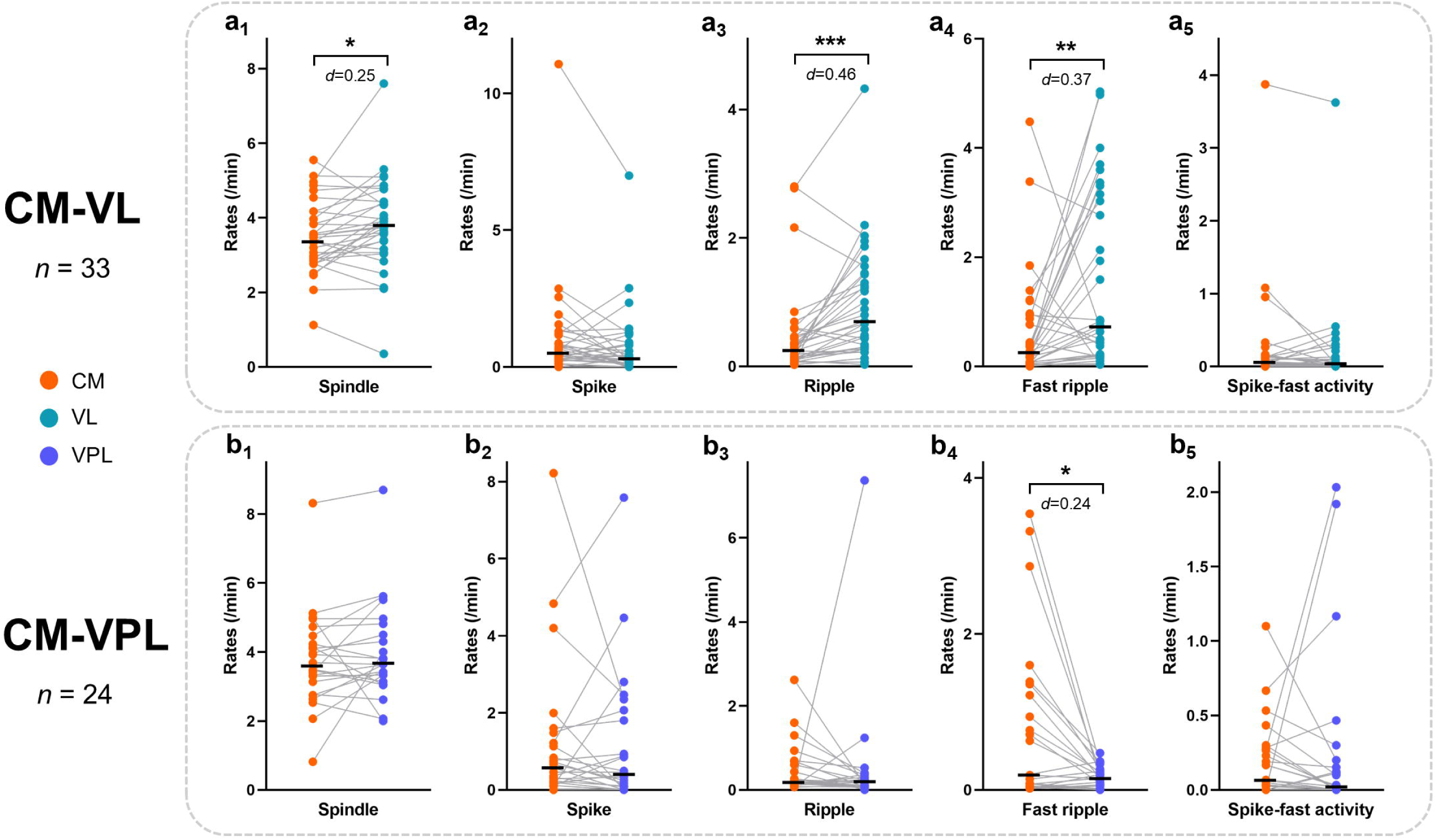
Pairwise comparisons of events’ rates between different nuclei. (a) The VL has more spindles, ripples, and fast ripples than the CM. (b) The CM has more fast ripples than the VPL. Black lines indicate median values. Each dot represents one subject. *, p < 0.05; **, p < 0.01; ***, p < 0.001; CM, centromedian nucleus; VL, ventral lateral nucleus; VPL, ventral posterolateral nucleus.

### 3.4 The frequency of thalamic spindles and HFOs

The median frequency of thalamic spindles was 11.91 Hz (IQR = 11.32–12.60 Hz) for the thalamus as a whole. Median spindle frequencies by nucleus were as follows: 11.72 Hz in the CM, 11.72 Hz in the VL, 12.50 Hz in the VPL, and 12.89 Hz in the Pu.

To further characterize spindle dynamics, we subdivided spindles into slow (10–12 Hz) and fast (12–16 Hz) spindles.^50,56^ Median rates of slow spindles (events per minute) were: 1.53 in the whole thalamus, 1.69 in the CM, 2.02 in the VL, 1.27 in the VPL, and 0.97 in the Pu. Median rates of fast spindles were: 1.67 in the whole thalamus, 1.33 in the CM, 1.76 in the VL, 1.96 in the VPL, and 4.66 in the Pu. Paired comparisons revealed that the Pu exhibited a significantly higher rate of fast spindles than slow spindles within individuals (*p* = 0.019, *d* = 0.73, *n* = 10, Fig. S8a). No significant differences between slow and fast spindle rates were observed in the other thalamic nuclei.

Frequency and duration of HFOs were also analyzed and summarized in Table S5. The median frequency of thalamic ripples was 124.37 Hz (IQR = 110.27–140.00 Hz), and the median frequency of fast ripples was 355.00 Hz (IQR = 311.25–381.25 Hz) (Fig. S8b). Pairwise comparisons revealed that ripple frequencies were significantly higher in the CM than in both the VL (*p* = 0.012, *d* = 0.34, *n* = 33) and the VPL (*p* = 0.005, *d* = 0.45, *n* = 30). In contrast, fast ripples in the VL had significantly higher frequencies than those in the CM (*p* < 0.001, *d* = 0.57, *n* = 23) (Fig. S8c).

### 3.5 Spike co-occurrences between the SOZ and thalamus

To quantify the co-occurrence of spikes between the SOZ and thalamus, we calculated the proportion of co-spikes using the number of co-occurring events as the numerator. Two adjusted percentages were derived by using either the total number of thalamic spikes or the total number of SOZ spikes as the denominator (**Fig. 2d**). The former reflects how thalamic spikes correlate with spikes in the SOZ, while the latter reflects the proportion of SOZ spikes that co-occur with thalamic spikes.

#### 3.5.1 Thalamic spikes do not always co-occur with SOZ spikes

When the thalamic spike count was used as the denominator, subjects with no thalamic spikes were excluded from the analysis. The median value of the adjusted percentage of co-spikes were as follows: 61.81% (Pu, IQR = 57.84%–65.78%), 24.12% (VPL, IQR = 17.54%–42.45%), 20.40% (whole thalamus, IQR = 8.26%–43.03%), 18.07% (CM, IQR = 4.46%–49.90%), and 13.14% (VL, IQR = 3.21%–28.44%). These findings suggest that, with the exception of the Pu, a considerable portion of thalamic spikes may occur independently of SOZ epileptiform discharges. This interpretation was further supported by the α-value obtained from surrogate analysis, which indicated that, in some patients, the observed co-spike counts did not significantly exceed chance levels (**Fig. 5a**). Furthermore, no significant difference in co-occurrence proportions was found between patients with favorable and unfavorable surgical outcomes. Similarly, no differences were observed between patients with SOZs located in different anatomical regions (Fig. S9).

**Figure 5.**
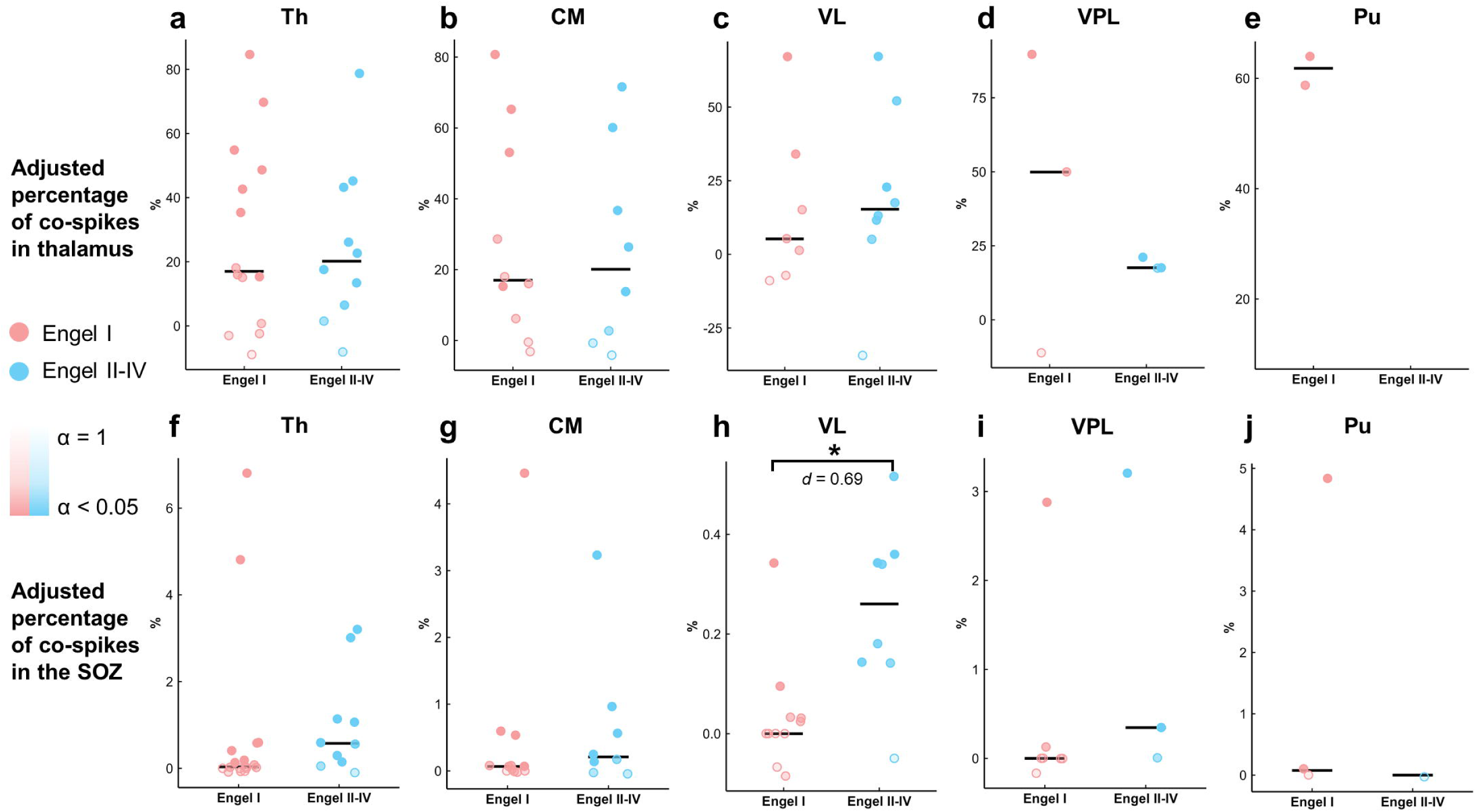
The proportions of co-spikes between the SOZ and thalamus in patients with focal SOZs and their relationships with surgical outcomes. (a)-(e) The percentage of co-spikes obtained when the number of thalamic spikes was used as the denominator. There was no difference between favorable and unfavorable surgical outcome group. (f)-(j) The percentage of co-spikes obtained when the number of SOZ spikes was used as the denominator. This proportion of co-spikes between the SOZ and VL was significantly higher in patients with unfavorable surgical outcomes. Each dot represents one subject. The transparency of the color indicates the alpha value in the surrogate data analysis. Lower transparency means that the number of co-spikes is higher than the chance level. SOZ, seizure onset zone; Th, thalamus; CM, centromedian nucleus; Pu, pulvinar nuclei; VL, ventral lateral nucleus; VPL, ventral posterolateral nucleus.

#### 3.5.2 SOZ spikes infrequently co-occur with thalamic spikes

When analyzing the spikes in the SOZ which co-occurred with thalamic spikes, median adjusted percentages of co-spikes were very low: 0.19% (IQR = 0.02%–0.69%) for the thalamus as a whole, 0.08% (IQR = 0.02%–0.69%) for CM, 0.03% (IQR = 0.02%–0.69%) for VL, 0.07% (IQR = 0.02%–0.69%) for VPL, and 0.04% (IQR = 0.02%–0.69%) for Pu. These results suggest that only approximately 1 in every 500 SOZ spikes co-occurred with a thalamic spike when random co-occurrences were excluded (Fig. S10). To explore the potential drivers of these co-occurrence patterns, we analyzed directed functional connectivity between the SOZ and thalamic nuclei. We found that the proportion of SOZ-thalamus co-spikes was positively correlated with the strength of the SOZ-to-thalamus driving force (quantified via ASI), a relationship that was statistically significant in the CM nucleus (*ρ* = 0.49, *p* = 0.021; **Fig. 6g**). Notably, the proportion of co-spikes between the SOZ and VL was significantly higher in patients with unfavorable surgical outcomes than in those with favorable outcomes (*p* = 0.013, *d* = 0.69; **Fig. 5h**). No significant differences were observed across SOZ anatomical locations (Fig. S9). Regarding the relationship with seizure dynamics, a trend toward a negative correlation was observed between the proportion of SOZ-thalamus co-spikes and the latency of thalamic ictal involvement (*ρ* = -0.36, *p* = 0.051, Fig. S6f), implying that stronger interictal coupling may be associated with earlier thalamic recruitment during seizures.

**Figure 6.**
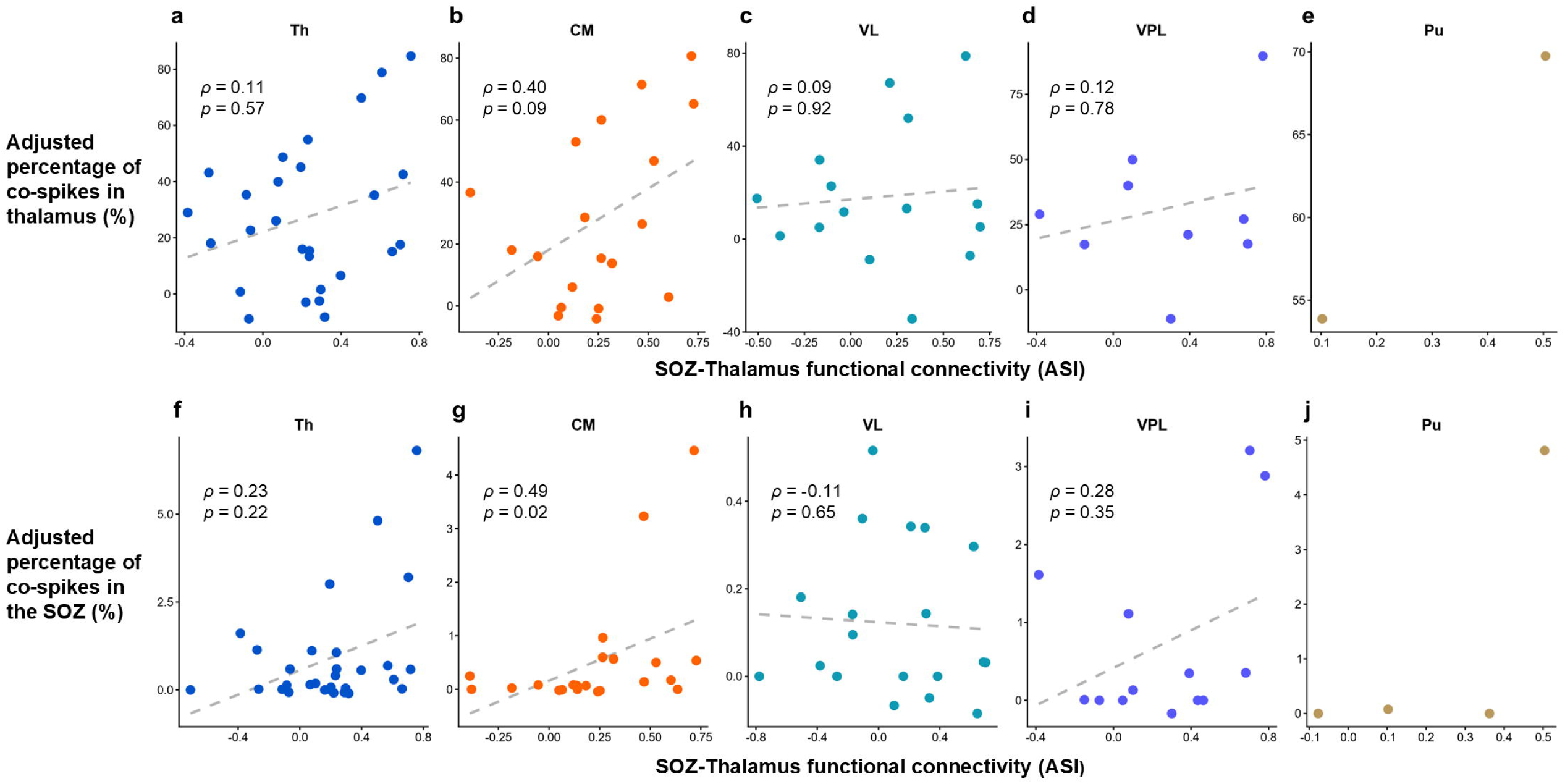
Correlation between SOZ-thalamus functional connectivity and spike co-occurrence in patients with focal SOZs. Directed functional connectivity was assessed using the Asymmetry Index (ASI) of Granger causality, where higher positive values indicate stronger drive from the SOZ to the thalamus. Spearman’s correlation coefficients (ρ) were calculated. (a)-(e) Correlation analysis where the percentage of co-spikes was calculated using the number of thalamic spikes as the denominator. (f)-(j) Correlation analysis where the percentage of co-spikes was calculated using the number of SOZ spikes as the denominator. A significant positive correlation was observed in the CM, indicating that stronger unidirectional functional connectivity from the SOZ to the CM is associated with a higher probability of spike co-occurrence. SOZ, seizure onset zone; Th, thalamus as a whole; CM, centromedian nucleus; Pu, pulvinar nuclei; VL, ventral lateral nucleus; VPL, ventral posterolateral nucleus.

#### 3.5.3 Pairwise comparisons between thalamic nuclei

No significant difference in the adjusted percentage of co-spikes was found between the CM and the VL, regardless of whether the spike count in the SOZ or the thalamus was used as the denominator (Fig. S11a). However, the CM exhibited a significantly higher co-occurrence percentage with SOZ spikes than the VPL when using the SOZ spike count as the denominator (*p* = 0.039, *d* = 0.47; Fig. S11b), indicating that SOZ spikes propagate more to the CM.

#### 3.5.4 Directionality of co-spikes between the SOZ and thalamus

In 74% of patients exhibiting co-occurring spikes between the SOZ and thalamus, the spike in the SOZ more likely preceded the spike in the thalamus (Fig. S12). In five patients, this directional pattern reached statistical significance, with all showing unidirectional propagation from the SOZ to thalamus. Analyses conducted separately for individual thalamic nuclei yielded similar results. These findings suggest that thalamic spikes typically follow those originating in the SOZ.

## 4 Discussion

Although thalamic implantation during SEEG is becoming increasingly common in the presurgical evaluation of drug-resistant epilepsy, the interictal electrophysiological signatures of the human thalamus remain poorly defined. This knowledge gap limits our ability to distinguish pathological from physiological activity and to harness thalamic recordings for surgical prognostication or neuromodulation programming. In this bicentric cohort of 64 patients, we provide the most comprehensive characterization of thalamic interictal events during NREM sleep to date. Our main findings were (1) thalamic SEEG events correlated with epilepsy prognosis: while SOZ-thalamus spike co-occurrence was rare, unfavorable outcomes were associated with elevated spike-fast activity, whereas sleep spindle rates within the CM and VL were reduced in surgically non-remediable patients; (2) event characteristics differed across nuclei, with the CM exhibiting stronger SOZ spike entrainment and more spike-fast ripples, whereas the VL exhibited higher spindle and HFO activity, underscoring the value of multisite thalamic recordings; and (3) the thalamus displayed unique electrophysiological patterns including spindle-coupled spikes, spike-like oscillations independent of cortical spikes, and physiological fast ripples, highlighting the complexity of thalamic SEEG review. Together, these results provide thalamic biomarkers of outcome while delineating physiological signatures essential for accurate SEEG interpretation and for refining thalamic neuromodulation strategies.

The overall rate of interictal epileptic events in the thalamus was low as demonstrated by this study, consistent with the traditional view that the thalamus is not highly epileptogenic. However, interictal epileptic events in the thalamus are still clinically important, reflecting thalamic involvement in the interictal epileptic network. Our results suggested that an extended interictal epileptic network, characterized by frequent thalamic spike-fast activity as well as a higher proportion of spikes in the SOZ propagating to the thalamus, could be a potential predictor of unfavorable seizure outcomes. In addition, thalamic spindle rates were reduced in patients deemed surgically non-remediable due to a more widespread epileptogenicity, possibly due to increased interictal spike-fast activity, as epileptiform discharges can disrupt spindle generation.^25^ These findings suggest the potential utility of thalamic interictal activity for guiding surgical decision-making.

Complementary findings in the Pu have been reported by Biagioni et al., who showed that patients with poor surgical outcomes (Engel III–IV) exhibited more thalamic spikes and fast ripples during NREM sleep.^27^ We only observed more spikes in surgically non-remediable patients than in those with favorable surgical outcomes (Engel I), while no significant group differences were found for fast ripples. The discrepancy may reflect differences in sampling of thalamic nuclei across studies or the limited specificity of spikes and fast ripples in characterizing thalamic epileptic activity.^29,44^ Spike-fast activity seems to be a better epileptic biomarker than pure spikes and HFOs within the thalamus according to our findings. However, whether thalamic biomarkers can add new and effective information for predicting surgical outcomes remains to be determined through further analysis in combination with cortical interictal markers.

Whether the thalamus can produce spikes independently of the cortex remains a topic of debate.^26,57^ Addressing this question is challenging due to the inevitable chance co-occurrence and the inherent spatial limitations of intracranial EEG sampling. In this study, we calibrated the random co-occurrence through a surrogate analysis and found that more than half of spikes in the CM, VL, and VPL did not co-occur with SOZ spikes in most patients. This aligns with prior research indicating that the thalamus can exhibit spikes independent of cortical sources.^26^ **However, this does not point to the intrinsic thalamic generation of interictal epileptiform discharges.** There are three possible explanations for isolated spikes in the thalamus during NREM sleep: (1) they are epileptiform discharges that propagated from EEG unsampled cortical areas; (2) they are actually sleep-related physiological events with spike-like morphology; (3) they are epileptiform discharges caused by the insult of the electrode implantation (less likely explanation).

Definitively identifying an isolated thalamic spike is technically difficult due to incomplete brain coverage in intracranial EEG. However, we still postulate that some thalamic spikes could be non-epileptic events based on the following reasons: (1) isolated spikes were also observed in seizure-free patients (**Fig. S13**), whose epileptogenic zones were likely well sampled; (2) in some patients, cortical spikes did not propagate to the thalamus, yet their thalamic spikes were still identified (**Figs. S13a, d-f**); (3) the recordings we selected were not from the acute post-implantation phase, and no evidence suggests that the thalamus is easily irritated by SEEG electrode implantation.^14^

This study found that SOZ spikes rarely propagated to the thalamus. It should be noted, however, that this co-occurrence ratio may be underestimated (Fig. S14). The thalamic spikes propagating from the cortex sometimes have a relatively longer duration and lower amplitude (**Figs. 1f, 2a**). Thus, they can deviate from the conventional definition of epileptic spikes in the cortex, potentially leading to their omission by detectors or exclusion during manual verification processes (Figs. S3, S14). Beyond these technical factors, our functional connectivity and spike timing analyses indicated that thalamic spikes are primarily driven by the cortex in focal epilepsy. We speculate that thalamic neuromodulation may exert its therapeutic effect by influencing cortico-thalamic-cortical feedback loop. Future direct stimulation experiments are needed to confirm if modulating this circuit dampens cortical epileptogenicity, subsequently reducing spike propagation from the SOZ to the thalamus. Finally, we did not investigate HFO co-occurrence between the cortex and the thalamus. Unlike spikes, HFOs are generally considered spatially confined events with limited capacity to propagate to distant brain regions, given their high-frequency and low-amplitude characteristics.^58^

In addition to possible non-epileptic spikes in the thalamus, several unique thalamic interictal events were found. While interictal spikes are generally thought to disrupt spindle generation,^25^ we identified instances where spikes and spindles co-occurred within the same thalamic channel (**Fig. 1f**), potentially due to convergence of propagated spikes and spindles.

Furthermore, the thalamus is different from the normal cortex in the occurrence rate of HFOs. Thalamic ripples are less frequent compared to most cortical regions.^48^ **Interestingly, rates of thalamic fast ripples are higher than in normal cortical regions.**^48^ Cortical fast ripples are almost always pathological, and therefore hardly ever occur in normal tissues.^43,45^

However, this may not be the case in the thalamus, where Daida et al. reported that physiological fast ripples could be present in the thalamus during NREM sleep, possibly in phase-amplitude coupling with sleep spindles.^29^ We also identified this phenomenon in this study (**Fig. S15**). These physiological fast ripples likely account for the relatively high thalamic fast ripple rates observed in this study. Findings from Kokkinos et al. have also revealed characteristic sawtooth delta events in the thalamus during REM sleep.^30^ Other studies have reported that the human thalamus, particularly the pulvinar, can exhibit specific delta band activity during REM sleep, dissociating from the concurrent cortical activation.^59–61^ Taken together, these results underscore the presence of thalamus-specific interictal EEG patterns that merit further exploration, particularly through visual observation of SEEG recordings.

This study is the first to systematically illustrate and compare the rates of classic interictal events across the CM, VL, and VPL. The CM showed higher rates of fast ripples and spike-fast ripples than the VPL, suggesting greater epileptic activity in this region. In contrast, the VL exhibited higher rates of sleep spindles and HFOs than the CM. Given that thalamic HFOs can be either physiological or pathological, this finding implies that the VL may engage in more physiological activity than the CM during NREM sleep. Furthermore, higher frequency ripples are generally considered more likely to be pathologic,^43^ and the CM exactly had a significantly higher frequency of ripples than the VL and VPL. Therefore, our findings indicate that the CM is possibly more epileptogenic in the interictal state.

The CM is a regulator of the cortico-thalamo-cortical network with widespread connections to the cortex, and is increasingly recognized as a key hub involved in both sleep regulation and epilepsy.^35^ Animal studies using passive recordings as well as optogenetic stimulation have demonstrated that the CM contributes to sleep–wake modulation.^62–64^ Neuromodulation targeting the CM has shown therapeutic benefits in patients with generalized and some types of focal epilepsy,^65–69^ which might be related to the role of the CM in sleep. A recent study reported that high-frequency CM stimulation during sleep disrupted sleep structure and exacerbated seizures, while low-frequency stimulation improved both sleep quality and seizure control.^70^ Our results verified that the CM participated in the interictal epileptic network during NREM sleep, especially in patients with unfavorable outcomes, providing insights into CM neuromodulation.

Moreover, our data suggest that VL SEEG recordings also yield valuable information for presurgical evaluation. We observed divergent factors influencing SOZ-thalamus spike co-occurrence: while the CM exhibited a significant positive correlation between functional connectivity strength and co-spike probability and other nuclei showed positive trends (implying a dependence on physiological pathway strength), the VL did not show this pattern. Instead, despite the low co-occurrence proportion of VL spikes with the SOZ, this proportion significantly stratified surgical outcomes. This dissociation implies that SOZ-VL co-occurrence is not merely a byproduct of baseline connectivity strength but may stem from a unique pathological feature related to epileptic networks. Thus, the present study highlights the importance of multi-nucleus thalamic recordings in understanding thalamic involvement in epilepsy. Future cohorts encompassing diverse epilepsy etiologies are needed to validate the distinct electrophysiological features of the VL.

Our study has some limitations. SEEG spatial sampling was restricted, as the coverage of the anterior thalamus was absent and the sample size for the Pu was small. The predominance of single-trajectory thalamic implantations, in order to reduce surgical risk, restricted intra-patient comparisons between various nuclei and may have reduced the statistical power to detect subtle differences, particularly given our conservative use of non-parametric tests. In addition, we did not have access to patient-specific Diffusion Tensor Imaging and histopathological confirmation for patients treated with laser ablation or neuromodulation, which precluded personalized analyses of structural connectivity and prevented stratification by etiology. Finally, since our analysis focused exclusively on NREM sleep, our findings may not be fully generalizable to wakefulness or REM sleep. The characterization of sleep-related non-epileptic spikes also requires further corroboration in future studies.

In conclusion, this study investigated interictal epileptic and non-epileptic transient events in the thalamus using multisite SEEG recordings, with comparisons made across different outcome groups. To our knowledge, this is the first study to quantitatively assess electrophysiological events in the human CM, VL, and VPL. This study enhances our understanding of the distinct electrophysiological profiles of thalamic nuclei and supports the use of thalamic recordings in presurgical evaluation. Further research is required to thoroughly elucidate the unique non-epileptic electrophysiological activity of the thalamus during sleep and wakefulness, thereby avoiding the misinterpretation of normal thalamic events as epileptiform.

## Contributors

**Hongyi Ye:** Writing – review & editing, Writing – original draft, Visualization, Validation, Software, Methodology, Investigation, Formal analysis, Data curation, Conceptualization. **Kassem Jaber:** Writing – review & editing, Visualization, Software, Resources, Data curation. **Alyssa Ho:** Data curation. **Lingqi Ye:** Writing – review & editing, Visualization, Software, Data curation. **John Thomas:** Writing – review & editing, Methodology. **Xiaowei Xu:** Writing – review & editing, Visualization. **Cong Chen:** Writing – review & editing, Data curation, Funding acquisition. **Yihe Chen:** Data curation. **Guoping Ren:** Data curation. **Matthew Moye:** Data curation. **Tamir Avigdor:** Writing – review & editing, Methodology. **Irena Dolezalova:** Methodology, Data curation. **Petr Klimes:** Writing – review & editing, Data curation. **Prachi Parikh:** Data curation. **Derek Southwell:** Writing – review & editing, Data curation. **Zhe Zheng:** Data curation. **Junming Zhu:** Data curation. **Shuang Wang:** Writing – review & editing, Validation, Supervision, Resources, Project administration, Funding acquisition, Conceptualization. **Birgit Frauscher:** Writing – review & editing, Writing – original draft, Validation, Supervision, Resources, Project administration, Methodology, Investigation, Funding acquisition, Formal analysis, Conceptualization.

## Acknowledgments

We would like to express our gratitude to the staff and EEG technicians at the Epilepsy Centers of SAHZU and DUMC. We would also like to thank all the members of the ANPHY lab for their suggestions on methodology and feedback on figures. This study was supported by start-up funding of Duke University to B.F., the National Natural Science Foundation of China (grant number: 82301636, 82471469), and the Zhejiang Provincial Natural Science Foundation (grant number: LD24H090003).

## Notes

### Competing Interest Statement

The authors have declared no competing interest.

### Summary of Updates

Manuscript, figures, and supplementary materials have been updated. A new author has been added to the authorship list.

