## Supplementary Materials for "Thalamic Interictal Epileptic and Non-Epileptic Events during NREM Sleep in Patients with Focal Epilepsy: a Stereo-EEG Study"

**Table S1. Demographic and clinical data of patients**

| Patient No. | Gender | Age, Y | Epilepsy duration, Y | Seizure onset zone | Sampled nuclei | Histopathology | Seizure outcome / Follow up, M |
| --- | --- | --- | --- | --- | --- | --- | --- |
| ZJUT001 | Female | 39.4 | 18.4 | Lt T, P, O | CM, VL, MD | Gliosis | Engel IIb / 35.5 |
| ZJUT002 | Male | 13.8 | 6.8 | Rt F | CM, VL | FCD IIb | Engel IIb / 20.2 |
| ZJUT003 | Male | 33.1 | 6.1 | Lt P | CM, VL | FCD IIb | Engel IIIa / 34.3 |
| ZJUT004 | Female | 26.3 | 13.3 | Lt T | CM, VL | FCD IIIa | Engel IIb / 35.0 |
| ZJUT005 | Female | 29.4 | 17.4 | Lt T | CM, VL | N/A | Engel IVb / 35.8 |
| ZJUT006 | Male | 31.7 | 10.7 | Rt I | CM, VL | Gliosis | Engel IVb / 34.1 |
| ZJUT007 | Male | 21.7 | 16.0 | Rt F | CM, VL | FCD Ib | Engel Ia / 33.2 |
| ZJUT008 | Male | 26.8 | 2.5 | Rt F | CM, VL | FCD IIa | Engel Ia / 33.4 |
| ZJUT009 | Male | 25.4 | 12.4 | Diffuse | CM, VL | N/A | Surgically non-remediable |
| ZJUT010 | Female | 7.2 | 2.2 | Diffuse | CM, VL, Pf | N/A | Surgically non-remediable |
| ZJUT011 | Female | 19.3 | 10.3 | Rt MTL | CM, VL | HS | Engel Ia / 32.0 |
| ZJUT012 | Male | 17.2 | 5.2 | Diffuse | CM, VL | N/A | Surgically non-remediable |
| ZJUT014 | Male | 10.5 | 1.8 | Rt F | Pu, VL | FCD IIa | Engel IVb / 26.7 |
| ZJUT015 | Female | 19.6 | 12.6 | Lt F | CM, VL | TSC | Engel Ia / 26.8 |
| ZJUT016 | Male | 9.3 | 3.3 | Lt MTL | CM, VL | Non-specific | Engel Ia / 29.4 |
| ZJUT017 | Male | 16.9 | 7.9 | Lt P | CM, VL | FCD I | Engel IVa / 29.0 |
| ZJUT018 | Male | 39.9 | 27.9 | Lt MTL; Rt MTL | CM, VPL | N/A | Surgically non-remediable |
| ZJUT019 | Female | 31.1 | 10.1 | Lt MTL | CM, VL | Gliosis | Engel Ia / 28.3 |
| ZJUT020 | Female | 26.9 | 25.2 | Rt MTL | CM, VL | Gliosis | Engel Ib / 26.7 |
| ZJUT021 | Male | 34.0 | 32.0 | Lt MTL | CM, VL, Pf | HS | Engel Ia / 26.7 |
| ZJUT022 | Male | 27.1 | 4.2 | Lt MTL; Rt MTL | CM, VL | N/A | Surgically non-remediable |
| ZJUT023 | Female | 18.9 | 10.9 | Diffuse | CM, VL | N/A | Surgically non-remediable |
| ZJUT024 | Male | 11.7 | 4.0 | Rt T | CM, VPL | Gliosis | Engel IVb / 26.9 |
| ZJUT025 | Female | 13.1 | 7.1 | Rt F | CM, VPL | Gliosis | Engel Ia / 21.6 |
| ZJUT026 | Male | 25.4 | 13.4 | Rt, T, P, O | CM, VPL | N/A | Engel IIIa / 24.5 |
| ZJUT027 | Male | 9.7 | 8.7 | Rt I | CM, VPL | FCD IIa | Engel IVb / 23.4 |
| ZJUT028 | Male | 15.8 | 8.8 | Rt T, P, O | CM, VPL, Pf | FCD Ia | Engel Ib / 24.1 |
| ZJUT030 | Female | 22.7 | 7.7 | Rt T, P, O | CM, VPL | FCD Ia | Engel IVb / 22.0 |
| ZJUT031 | Female | 33.1 | 15.1 | Lt F | CM, VL, VPL | FCD IIb | Engel Ia / 22.5 |
| ZJUT032 | Male | 24.7 | 7.7 | Rt T, P, O | CM, VL, VPL | Gliosis | Engel Ib / 21.1 |
| ZJUT033 | Male | 19.3 | 14.3 | Lt F | CM, VL, VPL | Gliosis | Engel IIb / 20.9 |
| ZJUT034 | Female | 18.4 | 5.4 | Lt I | CM, VPL | FCD II | Engel Ia / 20.6 |
| ZJUT035 | Male | 15.6 | 7.6 | Lt MTL, I | CM, VPL | Gliosis | Engel Ia / 19.7 |
| ZJUT037 | Male | 25.4 | 22.4 | Lt I | CM, VL | Gliosis | Engel IIIa / 19.5 |
| ZJUT038 | Female | 19.0 | 11.0 | Lt T, P, O | CM, VL, VPL | Non-specific | Engel IVb / 20.0 |
| ZJUT039 | Male | 20.7 | 7.7 | Lt T | CM, VL | Gliosis | Engel Ib / 19.3 |
| ZJUT040 | Female | 27.5 | 23.5 | Lt F, T | CM, VL | N/A | Surgically non-remediable |
| ZJUT041 | Female | 31.1 | 18.6 | Lt T, P, O | CM, VL | N/A | Surgically non-remediable |
| ZJUT042 | Male | 45.8 | 33.8 | Diffuse | CM, VL | N/A | Surgically non-remediable |
| ZJUT043 | Male | 29.2 | 10.2 | Lt T | VL, MD | N/A | Engel Ia / 18.5 |
| ZJUT044 | Male | 16.3 | 11.3 | Lt F | Pu, VPL | N/A | Engel Ia / 16.5 |
| ZJUT045 | Female | 11.6 | 6.6 | Rt P | CM, VPL | Tumor | Engel Ia / 15.4 |
| ZJUT046 | Female | 27.6 | 12.6 | Lt T, O | CM, VPL | Heterotopia | Engel IIb / 12.0 |
| ZJUT047 | Female | 25.4 | 24.4 | Rt P | VPL | N/A | N/A |
| ZJUT048 | Female | 33.6 | 9.6 | Lt MTL | CM, VL, VPL | Gliosis | Engel IIa / 15.1 |
| ZJUT049 | Female | 33.4 | 10.4 | Rt F | VPL | Gliosis | Engel IVa / 14.0 |
| ZJUT051 | Male | 31.2 | 10.2 | Rt MTL | CM, VPL | N/A | N/A |
| DEP00003 | Female | 23.0 | 18.0 | Lt T | CM, VL, VPL | Gliosis | Engel IIb / 22.0 |
| DEP00005 | Female | 34.0 | 28.0 | Rt F, T | CM, Pu, VPL | FCD Ib | Engel IIIa / 16.0 |
| DEP00010 | Female | 19.0 | 2.0 | Diffuse | Pu, VPL | N/A | Surgically non-remediable |
| DEP00011 | Male | 30.0 | 25.0 | Rt T | Pu, VL, VPL | Non-specific | Engel Ia / 18.0 |
| DEP00020 | Female | 23.0 | 17.0 | Lt O | Pu | N/A | Surgically non-remediable |
| DEP00032 | Male | 38.0 | 10.0 | Lt T | Pu | N/A | Surgically non-remediable |
| DEP00041 | Female | 59.0 | 49.0 | Lt MTL | CM, VPL | N/A | Engel Ia / 18.0 |
| DEP00042 | Male | 18.0 | 8.0 | Lt T | CM, VPL | Non-specific | Engel IVb / 16.0 |
| DEP00062 | Male | 16.0 | 13.0 | Lt T | CM, VL, LGN | Gliosis | Engel Ia / 12.0 |
| DEP00067 | Female | 28.0 | 19.0 | Lt F, T | Pu, VPL | N/A | Surgically non-remediable |
| DEP00068 | Male | 24.0 | 13.0 | Lt T | CM, VPL | FCD Ic | Engel IIb / 12.0 |
| DEP00069 | Female | 25.0 | 21.0 | Lt O | Pu | N/A | Engel IIb / 12.0 |
| DEP00070 | Female | 41.0 | 13.0 | Lt MTL | Pu | Gliosis | Engel Ia / 12.0 |
| DEP00095 | Female | 30.0 | 24.0 | Lt F, T | CM, VL, VPL | N/A | Surgically non-remediable |
| DEP00101 | Female | 46.0 | 33.0 | Rt F | VPL | N/A | N/A |
| DEP00105 | Male | 45.0 | 28.0 | Lt T; Rt T | CM, VPL | N/A | Surgically non-remediable |
| DEP00108 | Female | 44.0 | 29.0 | Lt MTL; Rt T | Pu | N/A | Surgically non-remediable |

Y, years; M, months; Rt, right; Lt, left; F, frontal lobe; I, insular; O, occipital lobe; P, parietal lobe; T, temporal lobe; MTL, mesial temporal lobe; CM, centromedian nucleus; VL, ventral lateral nucleus; VPL, ventral posterolateral nucleus; MD, mediodorsal nucleus; Pf, parafascicular nucleus; Pu, pulvinar; LGN, lateral geniculate nucleus; HS, hippocampal sclerosis; FCD, focal cortical dysplasia; N/A, not available; TSC, tuberous sclerosis.

**Table S2 Subregions of the sampled Pulvinar**

| **Patient No.** | **Subregions** |
| --- | --- |
| ZJUT014 | Medial Pulvinar |
| ZJUT044 | Anterior Pulvinar |
| DEP00005 | Anterior Pulvinar |
| DEP00010 | Medial and Inferior Pulvinar |
| DEP00011 | Anterior Pulvinar |
| DEP00020 | Medial Pulvinar |
| DEP00032 | Medial and Inferior Pulvinar |
| DEP00067 | Inferior Pulvinar |
| DEP00069 | Medial Pulvinar |
| DEP00070 | Medial Pulvinar |
| DEP00108 | Medial and Inferior Pulvinar |

**Table S3. Interictal event detectors**

| **Detectors** | **Links** | **References** |
| --- | --- | --- |
| Spindle | N/A | (Mölle et al., 2011; Schiller et al., 2025) |
| Spike | <https://github.com/EpiReC-ISARG/IED_detector> | (Janca et al., 2015) |
| Spike-gamma | <https://doi.org/10.5281/ZENODO.11237651> | (Thomas et al., 2022) |
| HFO | <https://mni-open-ieegatlas.research.mcgill.ca/> | (von Ellenrieder et al., 2012) |

**Table S4. Multivariate analysis of the correlation between thalamic spike-fast activity and demographic factors on seizure outcomes**

| **Nuclei** | **Comparisons** | **Variates** | **Odds Ratio** | **95% CI** | ***p* value** |
| --- | --- | --- | --- | --- | --- |
| **Th** | Engel I  vs.  Others | Age | 1.03 | 0.95 – 1.14 | 0.47 |
|  |  | Duration | 0.97 | 0.88 – 1.07 | 0.54 |
|  |  | Gender | 1.03 | 0.31 – 3.50 | 0.96 |
|  |  | Spike-fast activity | **1.57** | 1.21 – 2.13 | 0.002 |
|  | Engel I  vs.  Engel II-IV | Age | 1.02 | 0.92 – 1.12 | 0.72 |
|  |  | Duration | 0.96 | 0.87 – 1.07 | 0.51 |
|  |  | Gender | 1.03 | 0.28 – 3.73 | 0.97 |
|  |  | Spike-fast activity | **1.48** | 1.10 – 1.99 | 0.01 |
|  | Engel I  vs.  Surgically non-remediable | Age | 1.06 | 0.95 – 1.18 | 0.28 |
|  |  | Duration | 0.97 | 0.86 – 1.10 | 0.86 |
|  |  | Gender | 1.01 | 0.22 – 4.56 | 0.99 |
|  |  | Spike-fast activity | **1.77** | 1.19 – 2.64 | 0.005 |
| **CM** | Engel I  vs.  Others | Age | 1.05 | 0.95 – 1.17 | 0.33 |
|  |  | Duration | 1.00 | 0.88 – 1.12 | 0.94 |
|  |  | Gender | 1.15 | 0.32 – 4.14 | 0.83 |
|  |  | Spike-fast activity | **1.42** | 1.08 – 1.96 | 0.02 |
|  | Engel I  vs.  Engel II-IV | Age | 1.05 | 0.94 – 1.17 | 0.38 |
|  |  | Duration | 0.98 | 0.86 – 1.11 | 0.71 |
|  |  | Gender | 1.03 | 0.25 – 4.17 | 0.97 |
|  |  | Spike-fast activity | **1.47** | 1.06 – 2.04 | 0.02 |
|  | Engel I  vs.  Surgically non-remediable | Age | 1.06 | 0.95 – 1.17 | 0.40 |
|  |  | Duration | 1.02 | 0.88 – 1.12 | 0.79 |
|  |  | Gender | 1.30 | 0.32 – 4.14 | 0.75 |
|  |  | Spike-fast activity | 1.34 | 1.08 – 1.96 | 0.11 |
| **VL** | Engel I  vs.  Others | Age | 1.02 | 0.90 – 1.16 | 0.75 |
|  |  | Duration | 0.97 | 0.84 – 1.10 | 0.63 |
|  |  | Gender | 0.62 | 0.11 – 3.24 | 0.57 |
|  |  | Spike-fast activity | **1.73** | 1.22 – 2.63 | 0.004 |
|  | Engel I  vs.  Engel II-IV | Age | 1.02 | 0.90 – 1.16 | 0.73 |
|  |  | Duration | 0.95 | 0.83 – 1.09 | 0.49 |
|  |  | Gender | 0.65 | 0.11 – 3.72 | 0.63 |
|  |  | Spike-fast activity | **1.56** | 1.05 – 2.32 | 0.03 |
|  | Engel I  vs.  Surgically non-remediable | Age | 1.02 | 0.87 – 1.19 | 0.82 |
|  |  | Duration | 1.00 | 0.85 – 1.17 | 0.99 |
|  |  | Gender | 0.56 | 0.07 – 4.50 | 0.59 |
|  |  | Spike-fast activity | **2.17** | 1.25 – 3.77 | 0.006 |

This generalized linear model included spike-fast activity rate, age, biological sex, and epilepsy duration as variables. An odds ratio less than zero indicates that the outcome is more likely to be of Engel I. *n_total_* = 60; Th, thalamus as a whole; CM, centromedian nucleus; VL, ventral lateral nucleus. Others: Engel II-IV and surgically non-remediable groups; The unit for age and epilepsy duration: years; The unit for spike-fast activity: logarithmic occurrence rates (/min).

**Table S5. Median frequency and duration of thalamic ripples and fast ripples across different nuclei**

| **Nuclei** | **Ripple** | | **Fast ripple** | |
| --- | --- | --- | --- | --- |
|  | Frequency (Hz) | Duration (ms) | Frequency (Hz) | Duration (ms) |
| Entire thalamus | 124.37 | 62.00 | 355.00 | 26.00 |
| CM | 150.00 | 53.00 | 307.50 | 28.50 |
| VL | 112.50 | 69.87 | 381.25 | 25.00 |
| VPL | 110.00 | 62.75 | 394.37 | 24.50 |
| Pu | 115.00 | 66.75 | 355.00 | 26.50 |

CM, centromedian nucleus; VL, ventral lateral nucleus; VPL, ventral posterolateral nucleus; Pu, pulvinar.

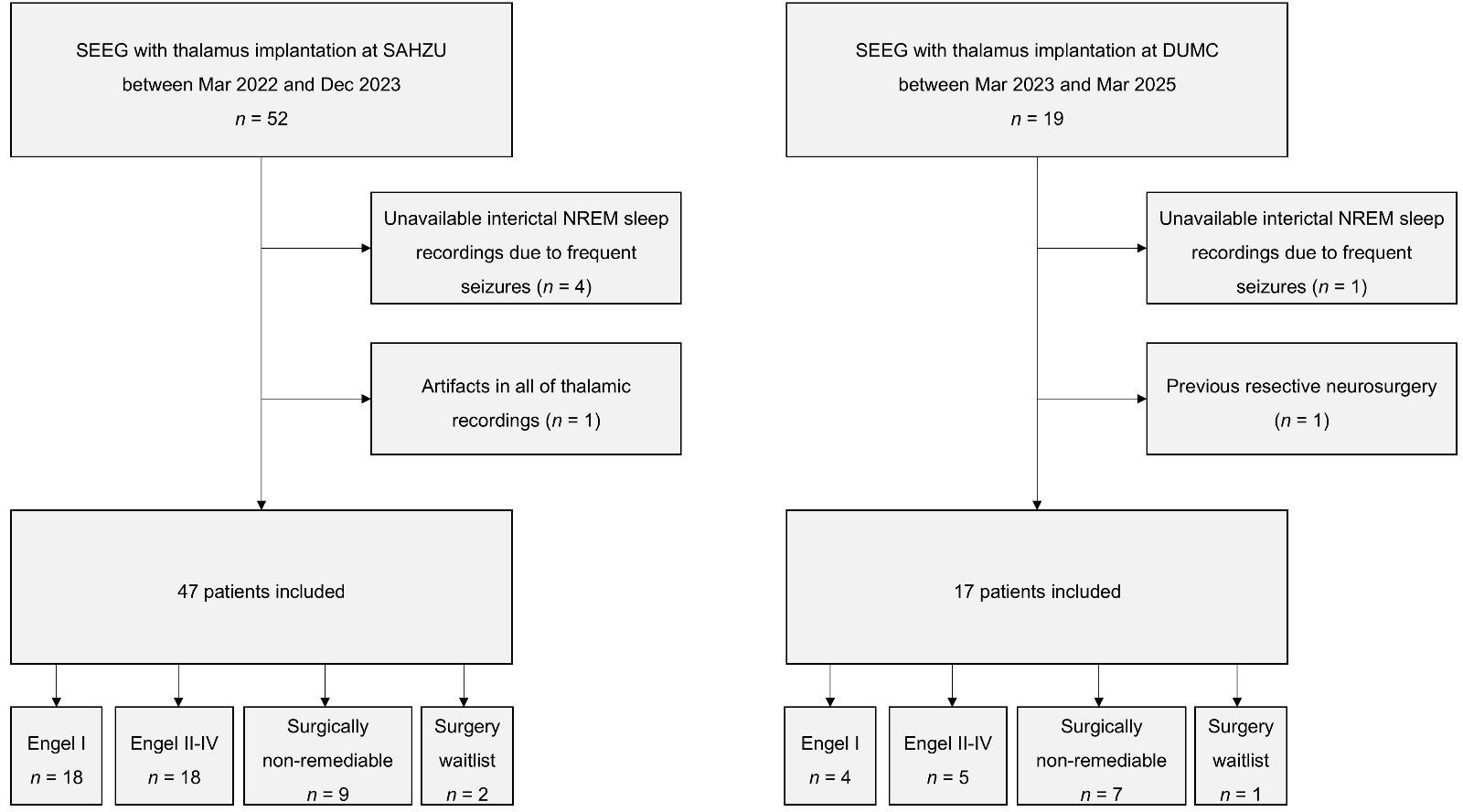

**Figure S1**. Patient inclusion and exclusion. Twenty-two, 23, and 16 patients were categorized into Engel I, Engel II-IV, and surgically non-remediable groups based on outcomes, respectively. SEEG, stereo-electroencephalography; SAHZU, The Second Affiliated Hospital Zhejiang University School of Medicine; DUMC, Duke University Medical Center; NREM, non-rapid eye movement.

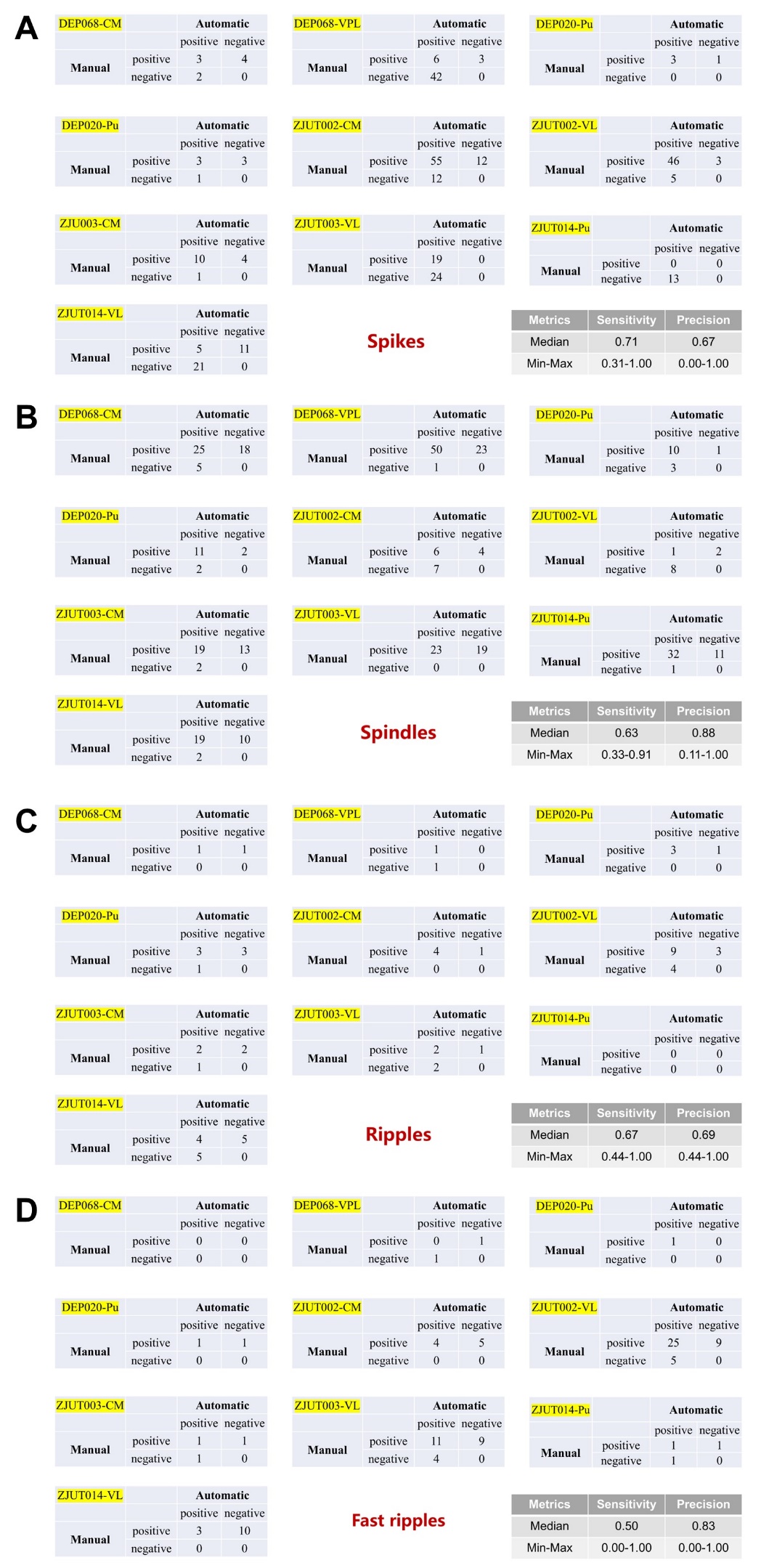

**Figure S2**. Performance of interictal transient event detectors based on 5-minute thalamic recordings. The automatically detected markers were labeled as either positive or negative according to the manual annotations. The upper left corner of the table shows the subject number and the thalamic nucleus in which the bipolar channel is located. Sensitivity and precision metrics are presented in the lower right corner of each figure. Overall, the detectors performed well. However, we observed that the automatic detection sometimes confused spikes and spindles, reflecting the need for visual verification. CM, centromedian nucleus; Pu, pulvinar nuclei; VL, ventral lateral nucleus; VPL, ventral posterolateral nucleus.

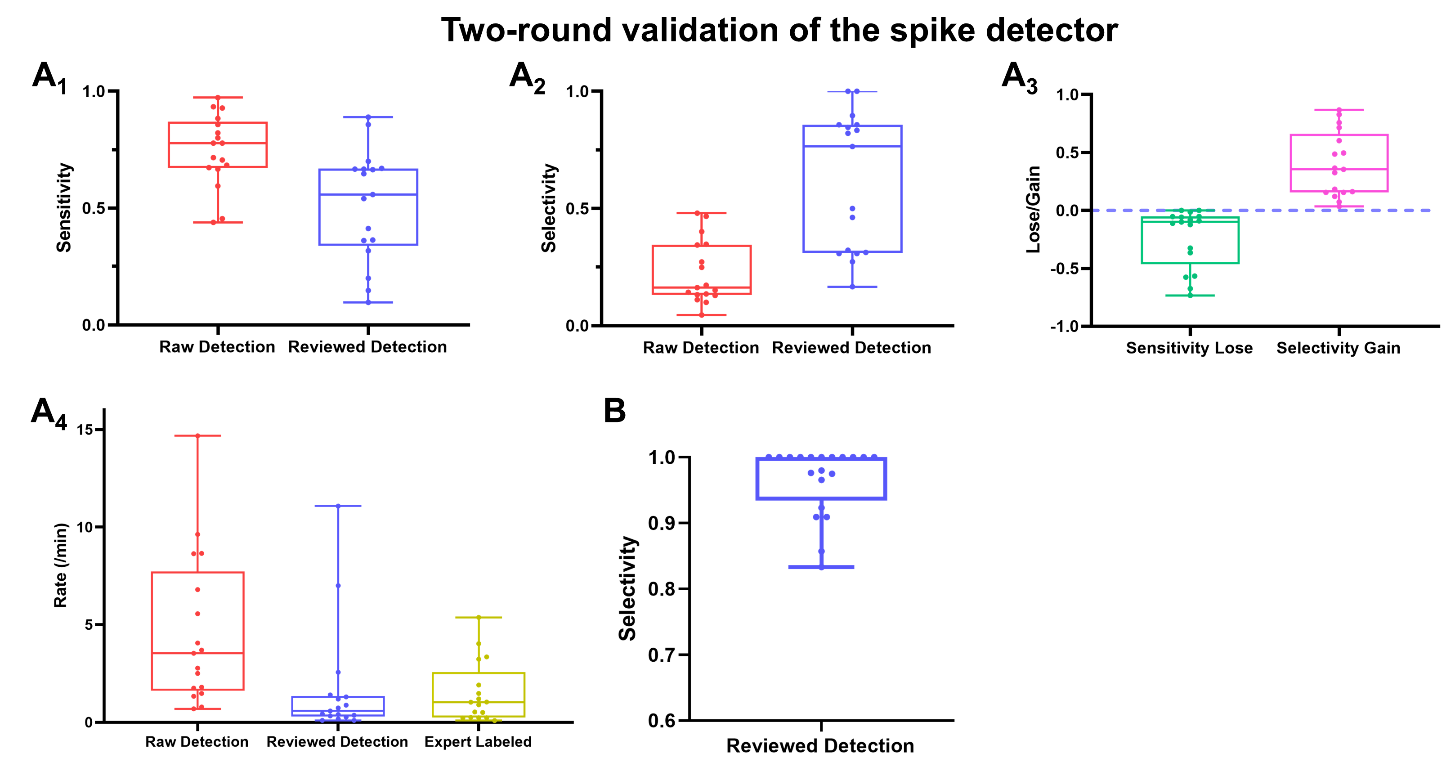

**Figure S3**. Validation of the semi-automated thalamic spike detection pipeline. To evaluate the trade-off between sensitivity and selectivity, a two-round validation was performed on a subset of ten patients (20 thalamic channels). A board-certified epileptologist (I.D.) manually annotated all spikes and sharp waves (20–200 ms) to serve as the Gold Standard. (A) Round 1: Comparison of detection rates. The boxplots display the event rates (events/min) detected by the raw automated detector, the semi-automated pipeline, and the Gold Standard. The raw detector yielded a significantly higher rate (median: 3.5/min, IQR: 1.7–6.8) compared to the Gold Standard (median: 1.0/min, IQR: 0.3–1.9), indicating a high false-positive rate likely due to physiological transients. The semi-automated pipeline yielded a more conservative rate (median: 0.6/min, IQR: 0.3–1.3), closer to the expert's assessment, reflecting reduced sensitivity but enhanced specificity. (B) Round 2: Evaluation of selectivity. The expert reviewed all events flagged by the semi-automated pipeline to determine precision. The results confirm high selectivity for the semi-automated method (median precision: 1.0, IQR: 0.95–1.0), ensuring that the detected events used for analysis were genuine epileptiform discharges.

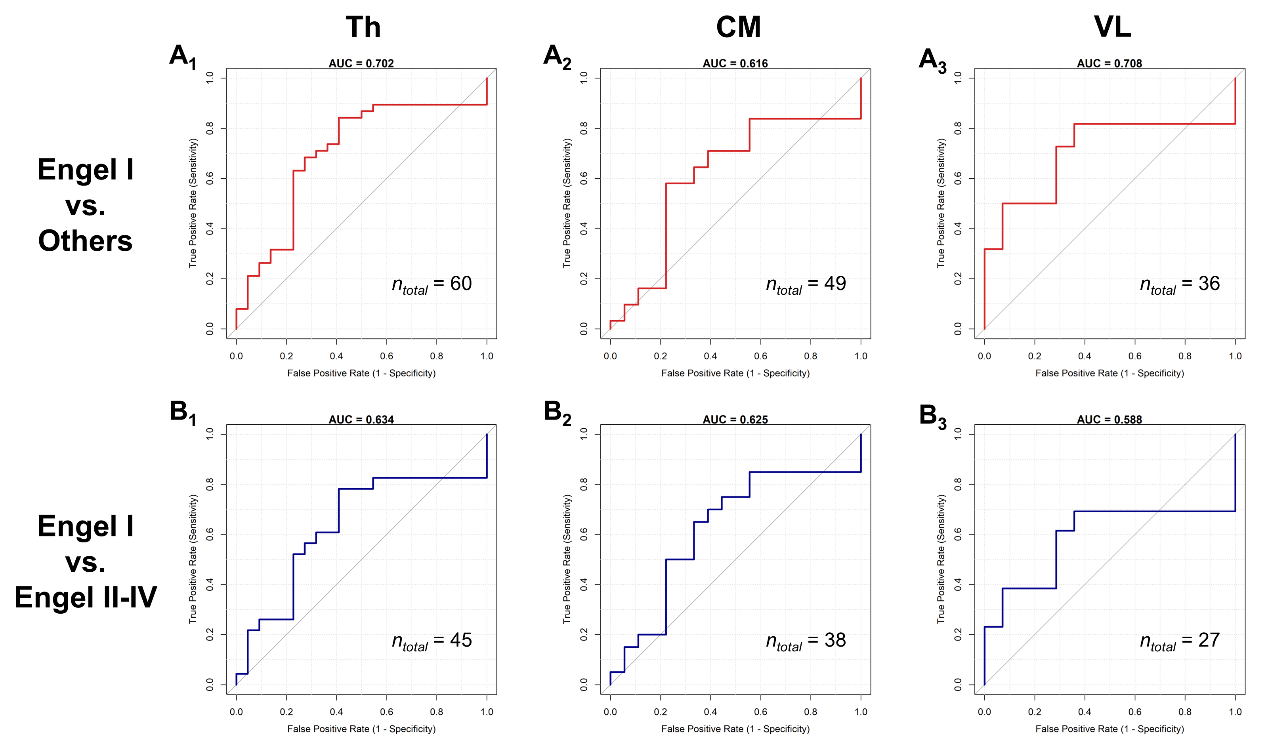

**Figure S4**. Predictive performance of thalamic spike-fast activity using Leave-One-Out Cross-Validation. (A) ROC curve distinguishing the Engel I group from others (the combination of Engel II-IV and surgically non-remediable groups). (B) ROC curve distinguishing the Engel I group from the Engel II-IV group. The Area Under the Curve (AUC) values are displayed within each plot. The unit for spike-fast activity: logarithmic occurrence rates (/min). ROC: Receiver Operating Characteristic.

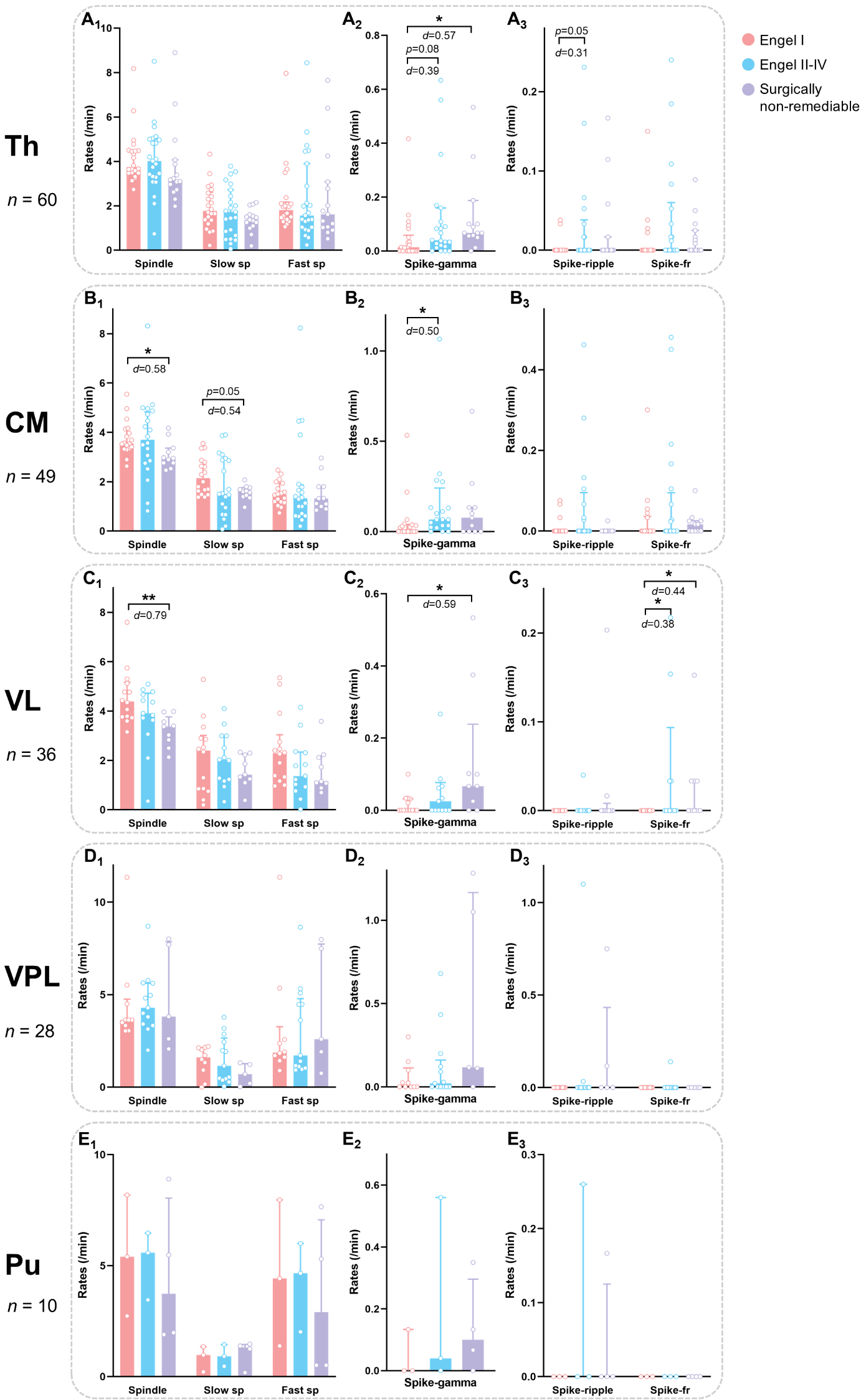

**Figure S5**. Rates of interictal transient events in different thalamic nuclei and in three outcome groups. (A) Event rates in the thalamus as a whole. Surgically non-remediable patients have more spike-gamma than patients with favorable surgical outcomes. (B) Event rates in the CM. Surgically non-remediable patients have fewer spindles than patients with favorable surgical outcomes. Patients with unfavorable surgical outcomes have more spike-gamma than those with favorable surgical outcomes. (C) Event rates in the VL. Surgically non-remediable patients have more spike-gamma, more spike-fast ripples, but fewer spindles than patients with favorable surgical outcomes. Patients with unfavorable surgical outcomes have more spike-fast ripples than those with favorable surgical outcomes. (D)-(E) There was no difference between three groups in the VPL and the Pu. Median values and interquartile ranges are shown in the plots. Each dot represents one subject. *, p < 0.05; **, p < 0.01; Th, thalamus; CM, centromedian nucleus; Pu, pulvinar nuclei; VL, ventral lateral nucleus; VPL, ventral posterolateral nucleus; sp, spindle; fr, fast ripple.

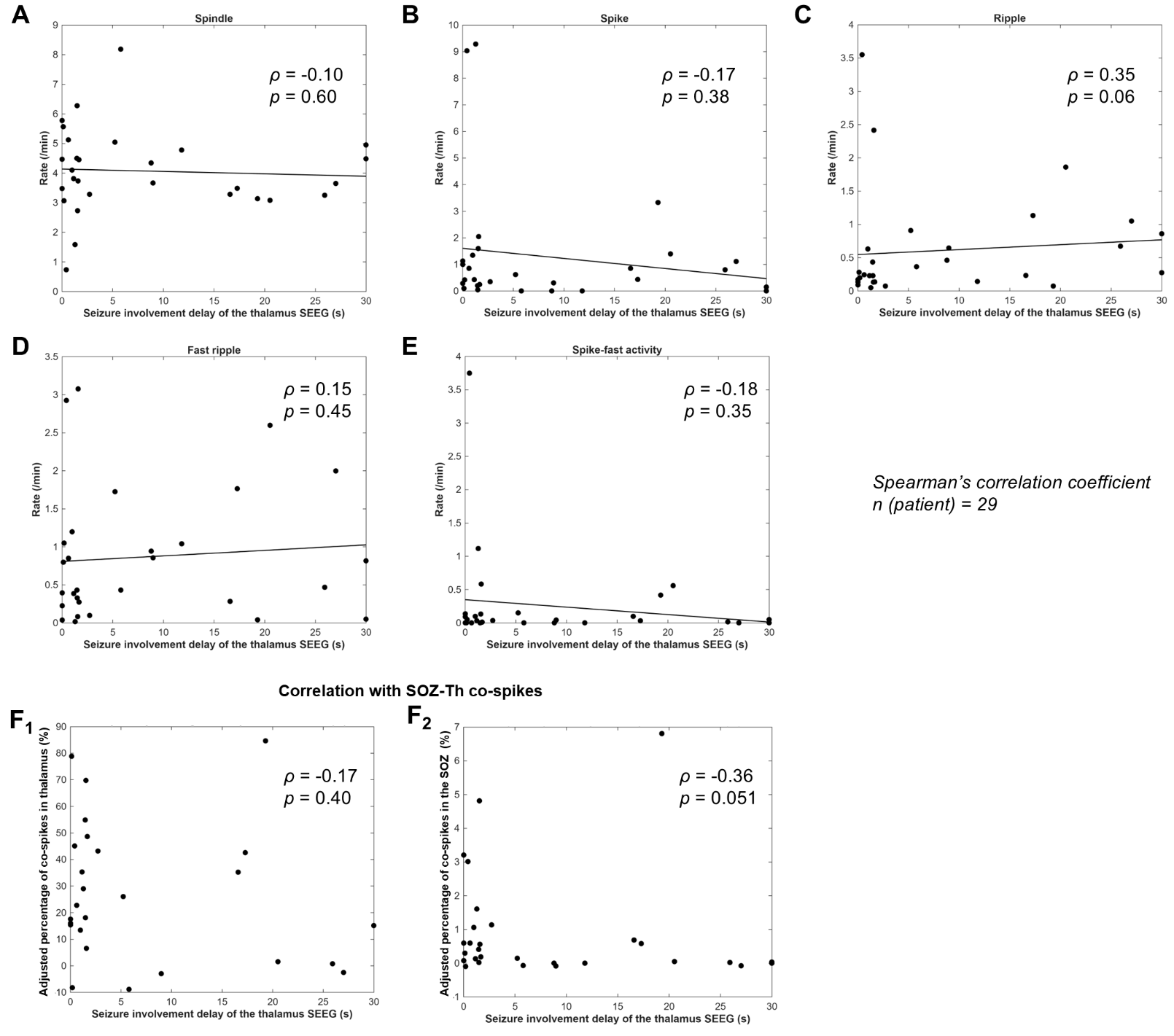

**Figure S6**. Correlation between thalamic ictal involvement and interictal metrics. The analysis was performed on a subset of 29 patients with focal SOZ. The X-axis represents the median delay of thalamic seizure involvement relative to the SOZ; cases with no involvement or delay >30s were censored at 30s. ρ represents the Spearman’s correlation coefficient. (A)-(E) Scatter plots showing the correlation between thalamic involvement delay and the rates of interictal events. No statistically significant correlations were found. There was a trend toward a positive correlation between the ictal involvement delay and thalamic ripple rates. (F) Correlation between thalamic ictal involvement delay and the adjusted proportion of SOZ-thalamic co-spikes. There was a trend toward a negative correlation between the delay and the proportion of SOZ-thalamic co-spikes, implying that shorter latency to thalamic involvement might associate with higher SOZ-thalamic co-spike occurrence. SOZ, seizure onset zone; Th, thalamus as a whole; SEEG, stereo-electroencephalography.

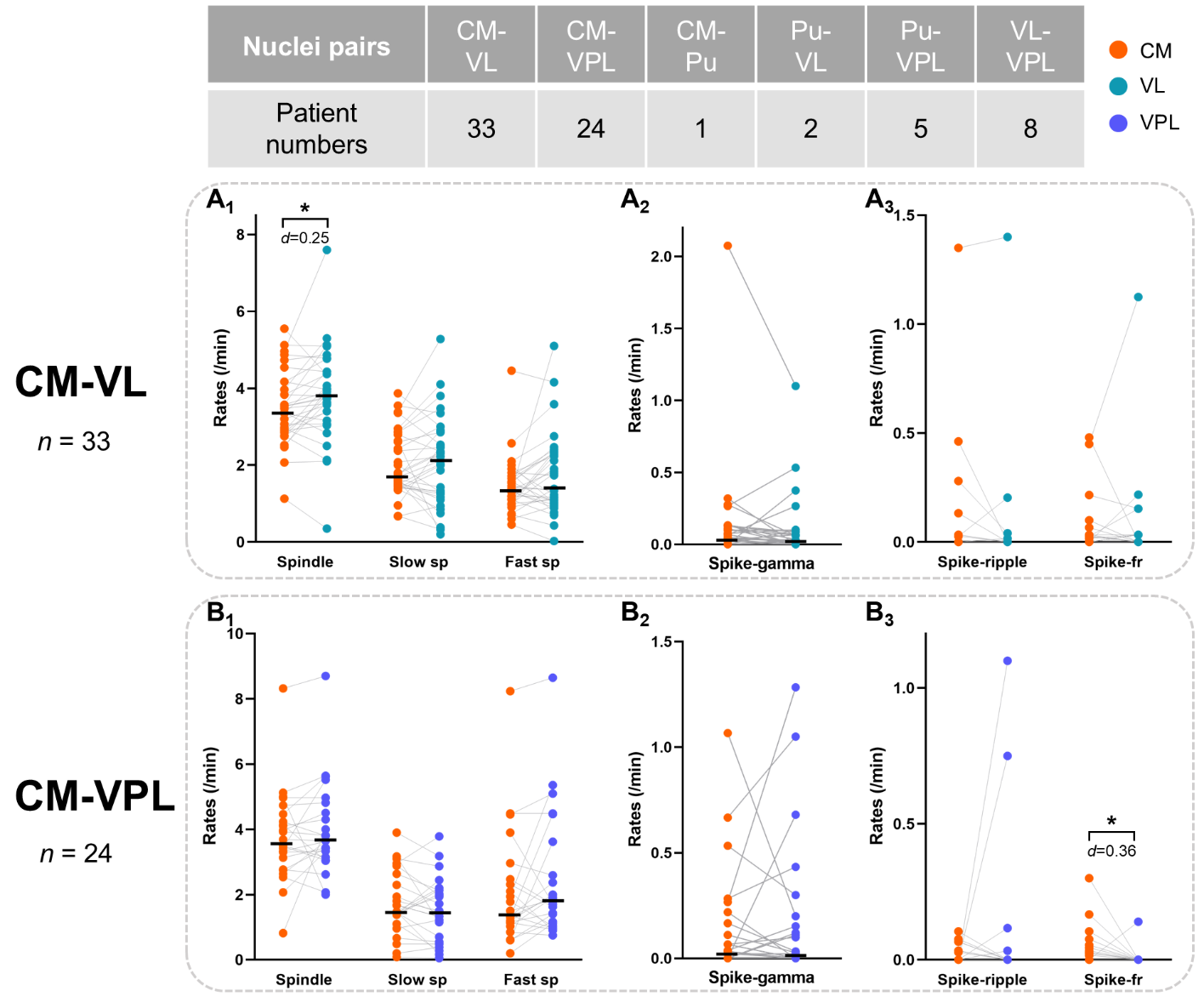

**Figure S7**. Pairwise comparisons of events’ rates between different nuclei. The table summarize the numbers of all the nuclei pairs within a subject. (A) The VL has more spindles than the CM, but there was no difference in the rates of slow and fast spindles between the VL and the CM. (B) The CM has more spike-fast ripples than the VPL. Black lines indicate median values. Each dot represents one subject. *, p < 0.05; CM, centromedian nucleus; Pu, pulvinar nuclei; VL, ventral lateral nucleus; VPL, ventral posterolateral nucleus; sp, spindle; fr, fast ripple.

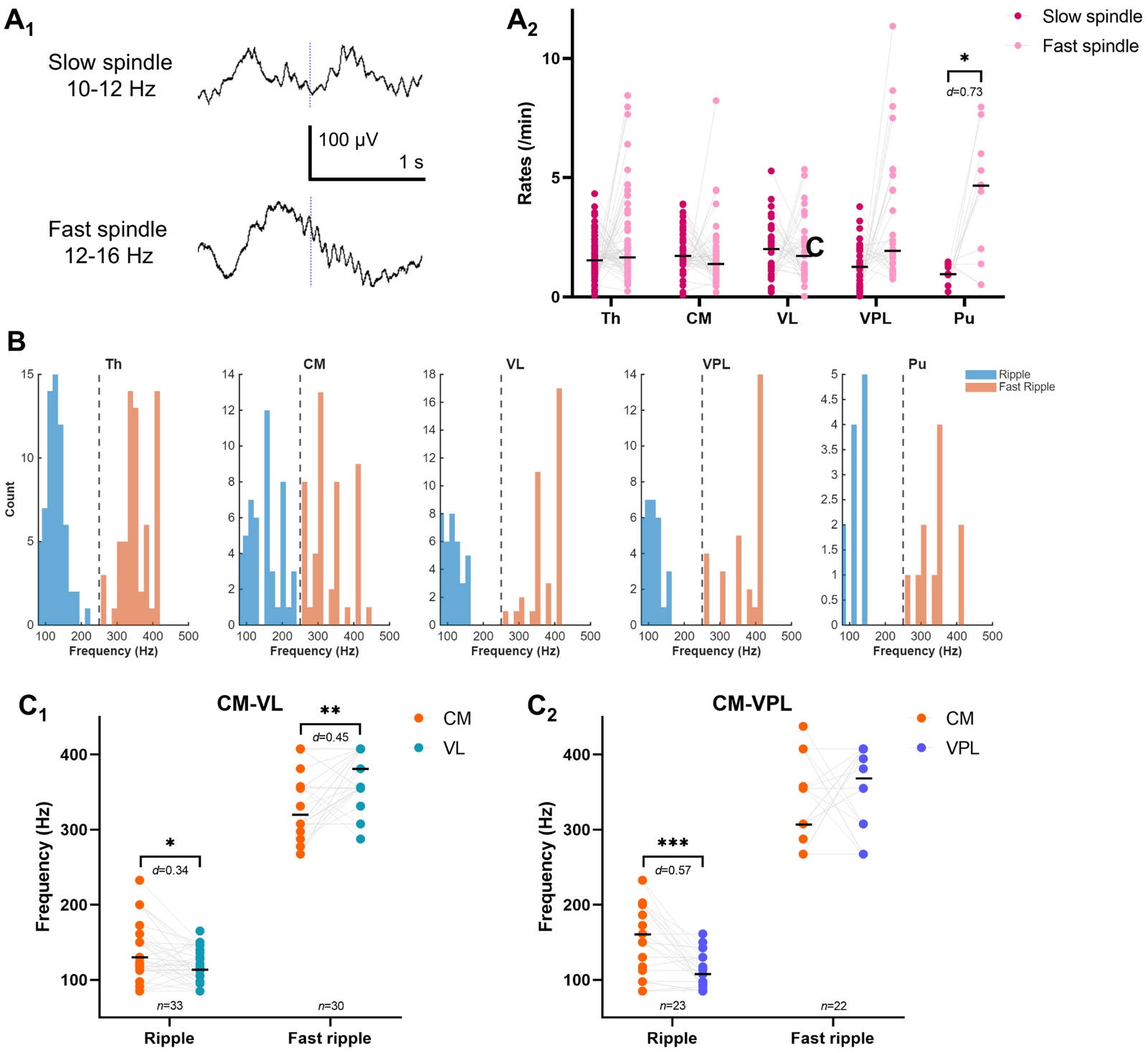

**Figure S8**. Frequency of spindles and HFOs across different thalamic nuclei. (A) Examples of slow spindles (10-12 Hz) and fast spindles (12-16 Hz), and the paired comparisons of their rates within individuals. We found that the Pu exhibited a significantly higher rate of fast spindles than slow spindles (*p* = 0.019, *d* = 0.73, *n* = 10). (B): Frequency distribution of thalamic HFOs. The histogram displays the aggregate frequency distribution of ripple and fast ripple events across the patient cohort. The y-axis represents the number of cases, and the gray dashed line indicates the cutoff frequency of 250 Hz. The distribution exhibits a bimodal structure. The ripple population is concentrated around 120 Hz, while the fast ripple population is primarily distributed between 300 and 400 Hz. Although a spectral shift was observed in the CM nucleus (with ripples trending towards higher frequencies and fast ripples towards lower frequencies), the two peaks remained distinct, supporting the use of the 250 Hz threshold. (C) Pairwise comparisons of the median frequency of ripples (80-250 Hz) and fast ripples (250-500 Hz) between different thalamic nuclei within individuals. Ripples in the CM had a higher frequency than ripples in the VL and VPL (*p* = 0.012, *d* = 0.34, *n* = 33; *p* = 0.0054, *d* = 0.45, *n* = 30), while fast ripples in the VL had a higher frequency than those in the CM (*p* = 0.00092, *d* = 0.57, *n* = 23). Black lines indicate median values. Each dot represents one subject. *, p < 0.05; **, p < 0.01; ***, p < 0.001; HFOs, high frequency oscillations; Th, thalamus; CM, centromedian nucleus; Pu, pulvinar nuclei; VL, ventral lateral nucleus; VPL, ventral posterolateral nucleus.

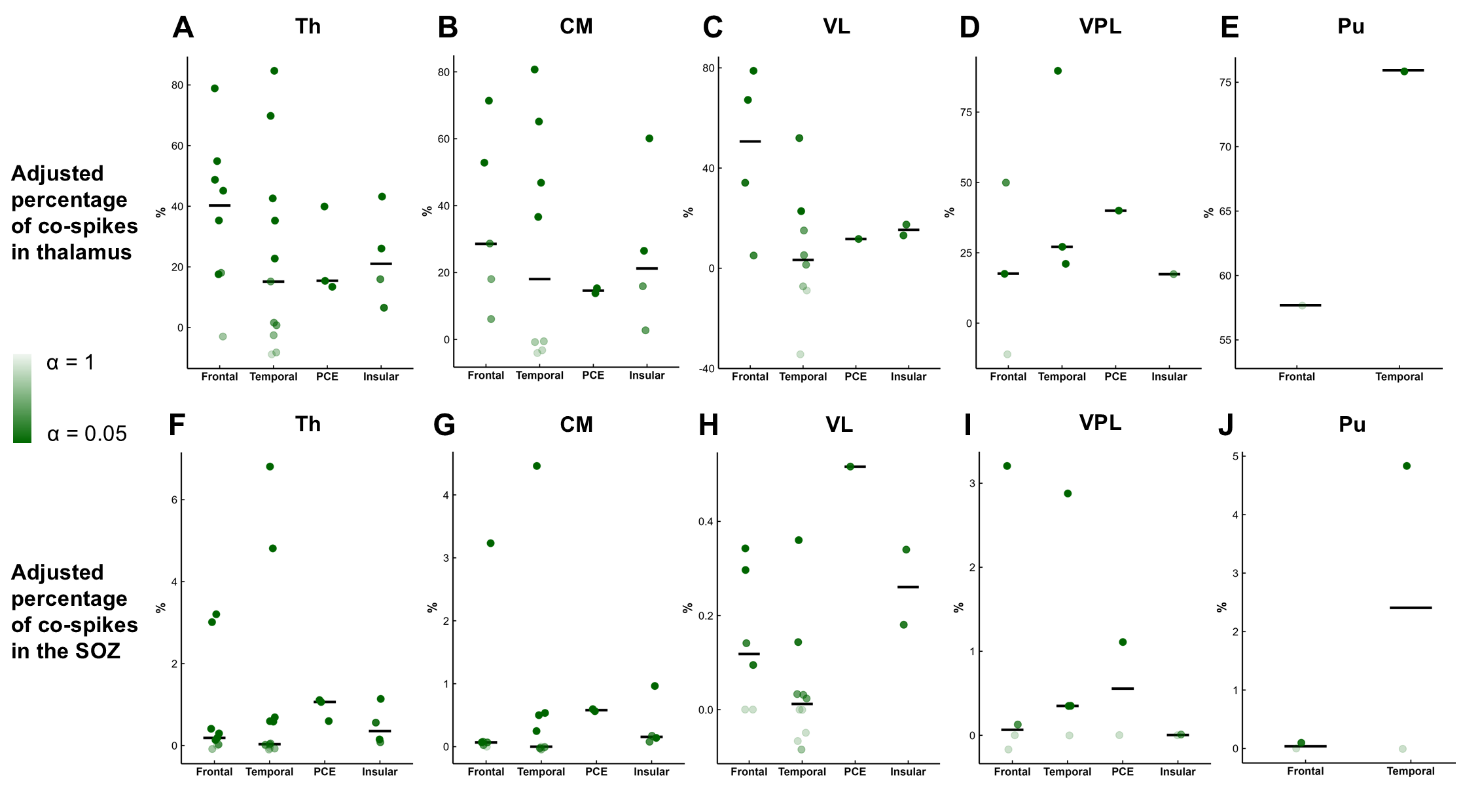

**Figure S9**. Proportion of co-spikes between the SOZ and the thalamus in different epilepsy types. The x-axis represents the type of epilepsy categorized according to the location of the SOZ. No statistically significant difference was observed between patients with SOZs located in different anatomical regions. Each dot represents one subject. The transparency of the dot color indicates the alpha value in the surrogate data analysis. Lower transparency indicates that the number of co-spikes is higher than the chance level. SOZ, seizure onset zone; PCE, posterior cortex epilepsy. Th, thalamus; CM, centromedian nucleus; Pu, pulvinar nuclei; VL, ventral lateral nucleus; VPL, ventral posterolateral nucleus.

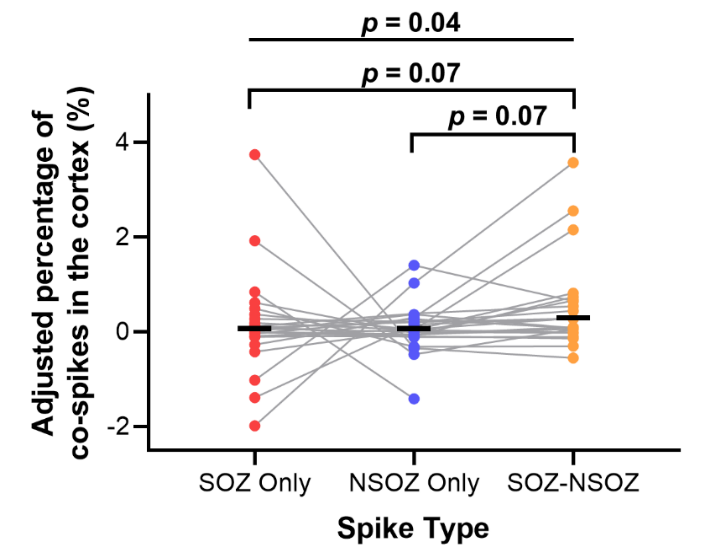

**Figure S10**. The adjusted proportion of co-spikes between cortical and thalamic spikes. Cortical spikes were categorized into three groups: SOZ-only (spikes confined to the SOZ), NSOZ-only (spikes confined to the NSOZ), and SOZ-NSOZ (synchronous spikes involving both zones). Adjusted percentages of co-spikes were derived by using the number of cortical spikes as the denominator. SOZ-NSOZ spikes exhibited a trend toward higher thalamic co-occurrence compared to focal spikes limited to the SOZ or NSOZ. Notably, the median adjusted cortical-thalamic co-spike proportion for the SOZ-NSOZ group was 0.27%, whereas they were nearly zero for the SOZ-only and NSOZ-only groups. This indicates that while widespread cortical spikes are more likely to involve the thalamus, the overall co-occurrence remains low.

Statistical test: non-parametric Friedman Test with post-hoc comparisons and Bonferroni correction. SOZ, seizure onset zone; NSOZ, non-SOZ.

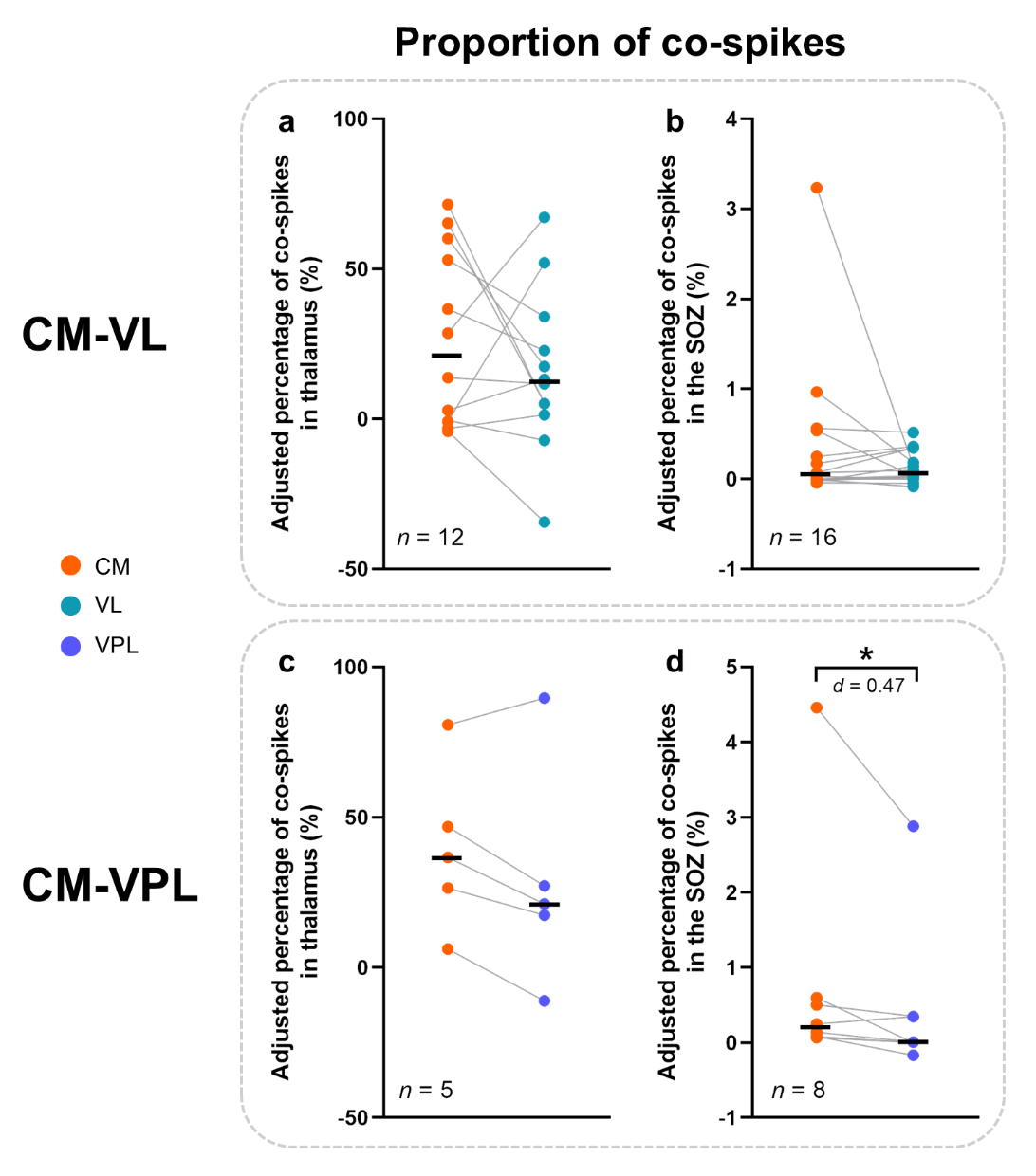

**Figure S11**. Pairwise comparisons of proportions of co-spikes with the SOZ between different nuclei. Percentages of co-spikes were calculated using the number of spikes in the thalamic nuclei and the SOZ as the denominator, respectively. (a)-(b) There was no significant difference between the CM and the VL. (c)-(d) Comparisons between the CM and the VPL demonstrated that spikes in the SOZ are more likely to propagate to the CM. Black lines indicate median values. Each dot represents one subject. *, p < 0.05; CM, centromedian nucleus; VL, ventral lateral nucleus; VPL, ventral posterolateral nucleus.

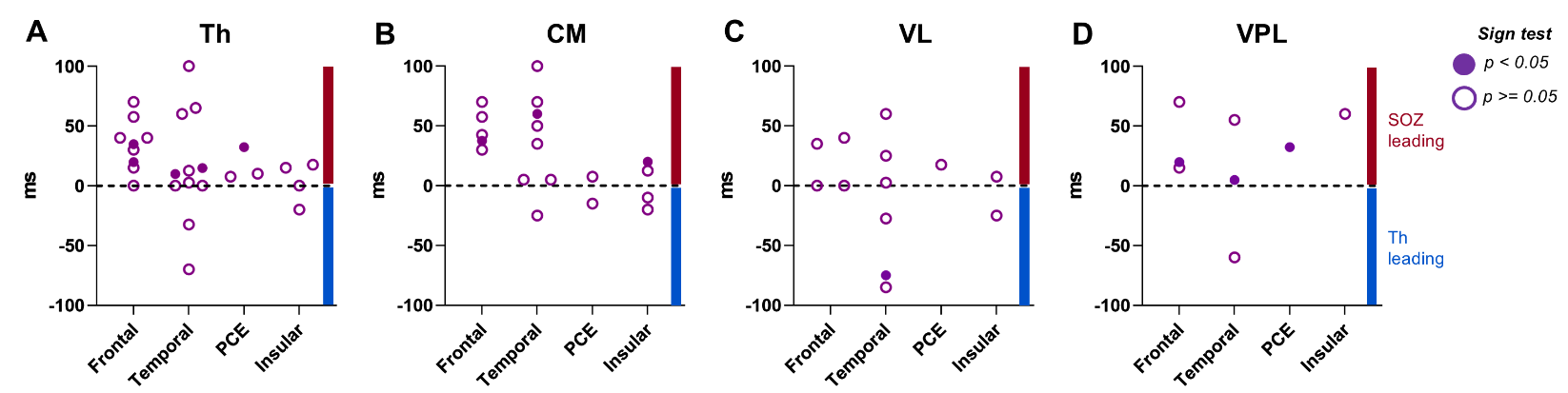

**Figure S12**. The median co-occurrence interval between the seizure onset zone and thalamic spikes. The x-axis represents the type of epilepsy categorized according to the location of the seizure onset zone. It shows that SOZ spikes more likely precede thalamic spikes. In five patients, this directional pattern reached statistical significance, with all showing propagation from the SOZ to the thalamus. Each dot represents one subject. The solid dots indicate significant unidirectional spike propagation between the seizure onset zone and the thalamic channel. PCE, posterior cortex epilepsy; Th, thalamus; CM, centromedian nucleus; Pu, pulvinar nuclei; VL, ventral lateral nucleus; VPL, ventral posterolateral nucleus.

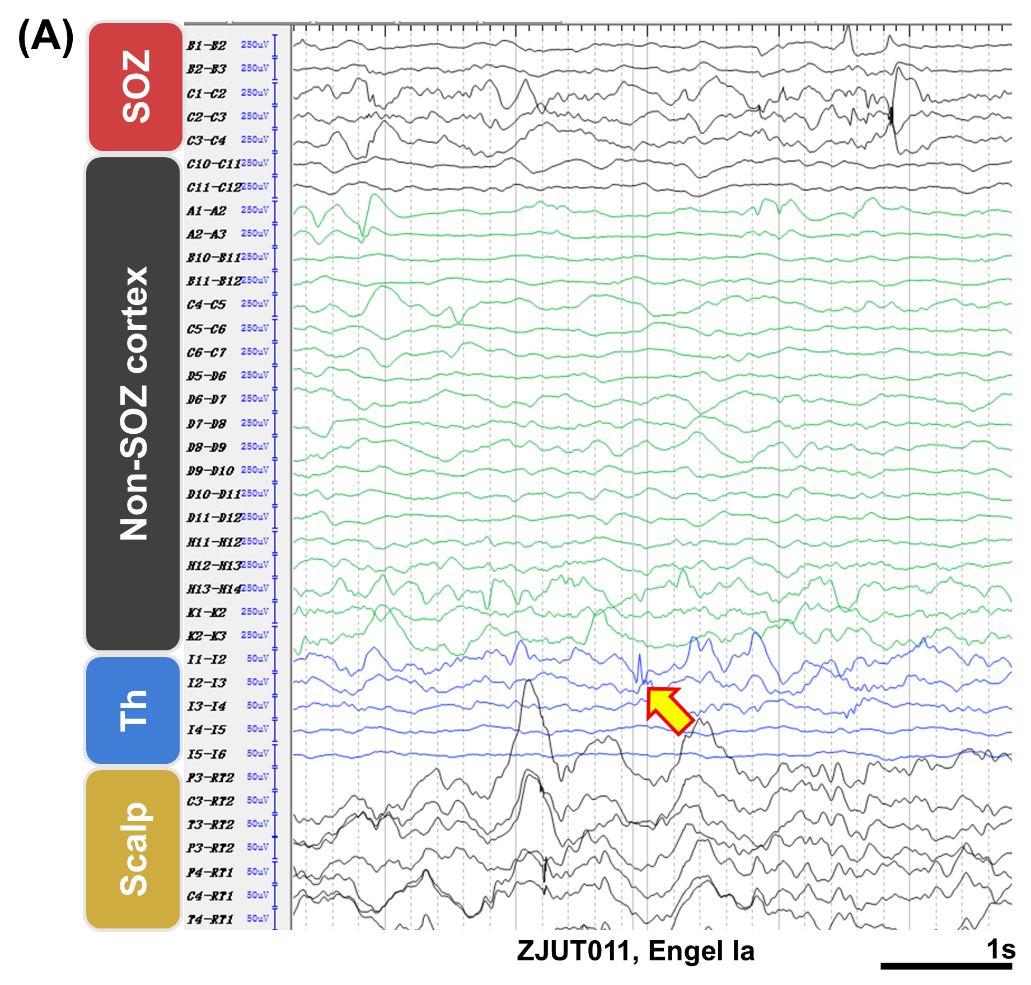

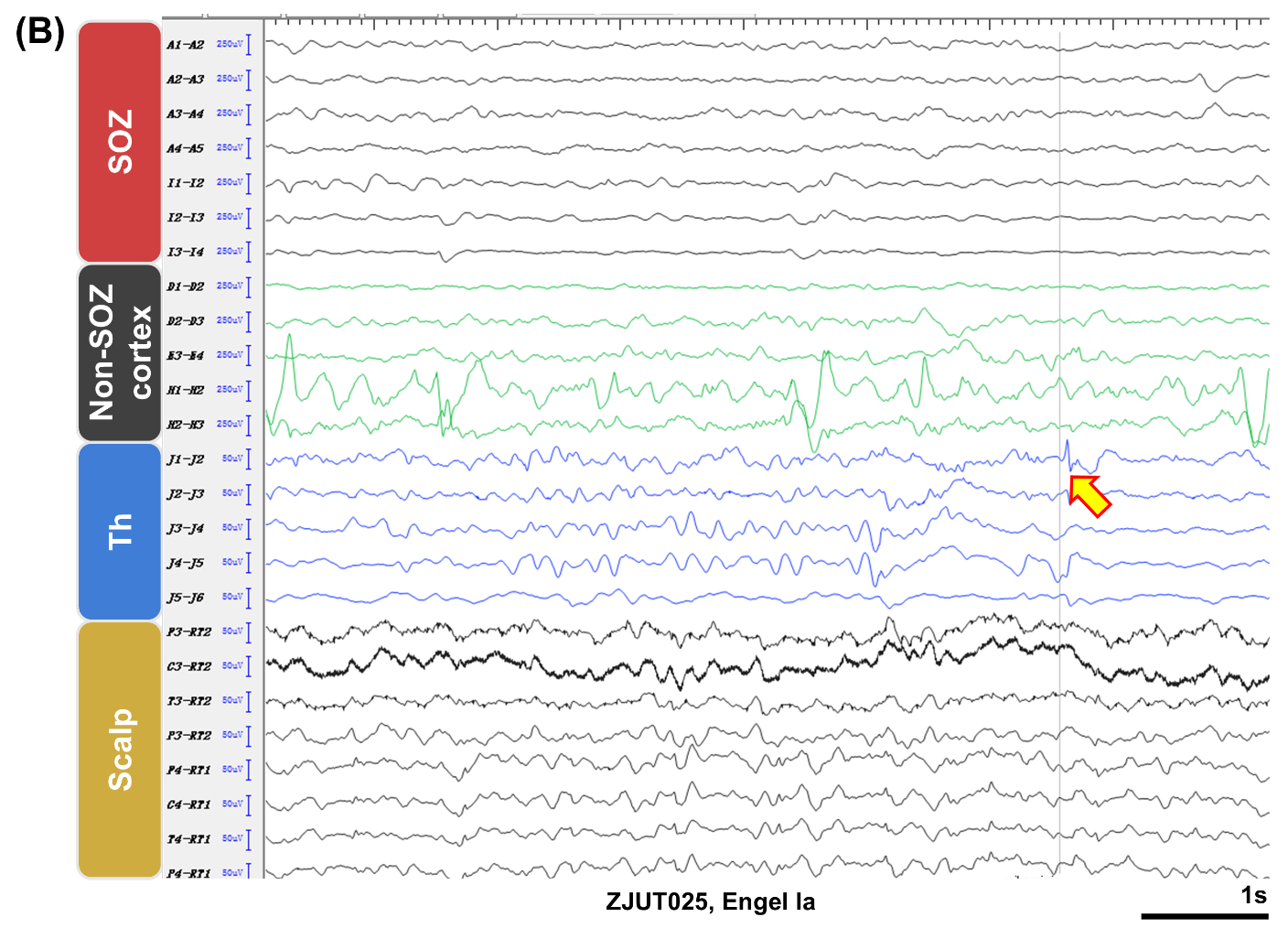

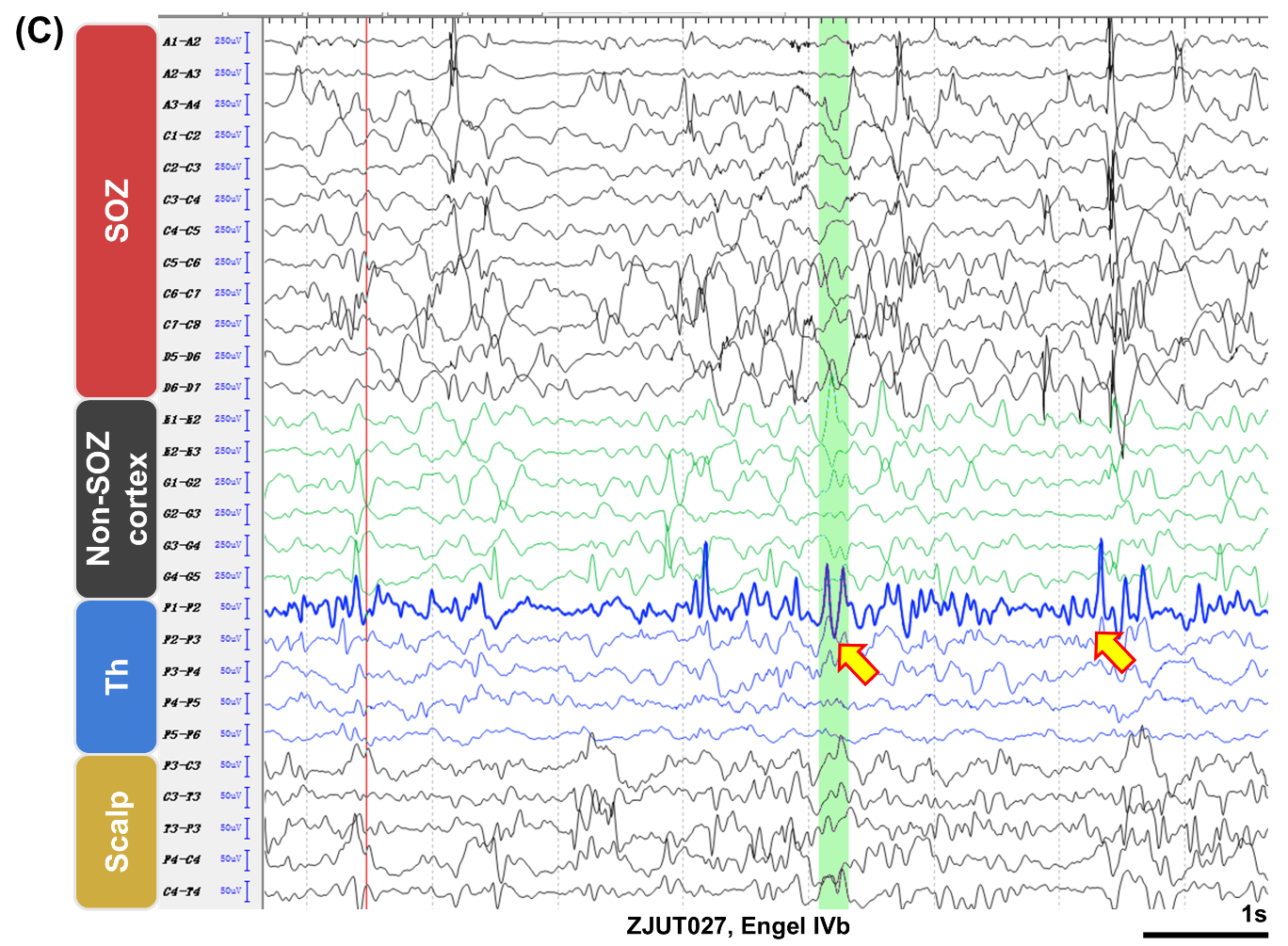

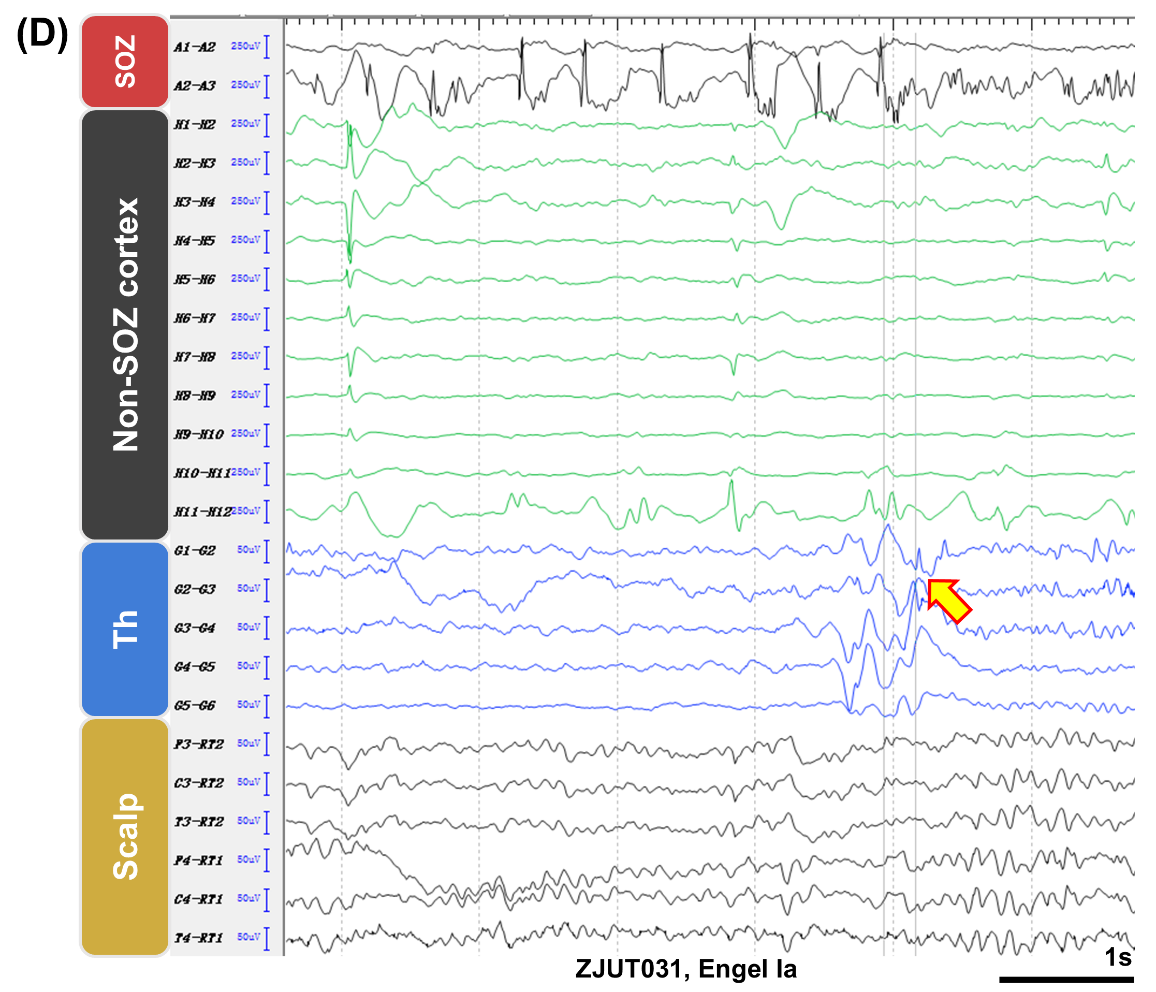

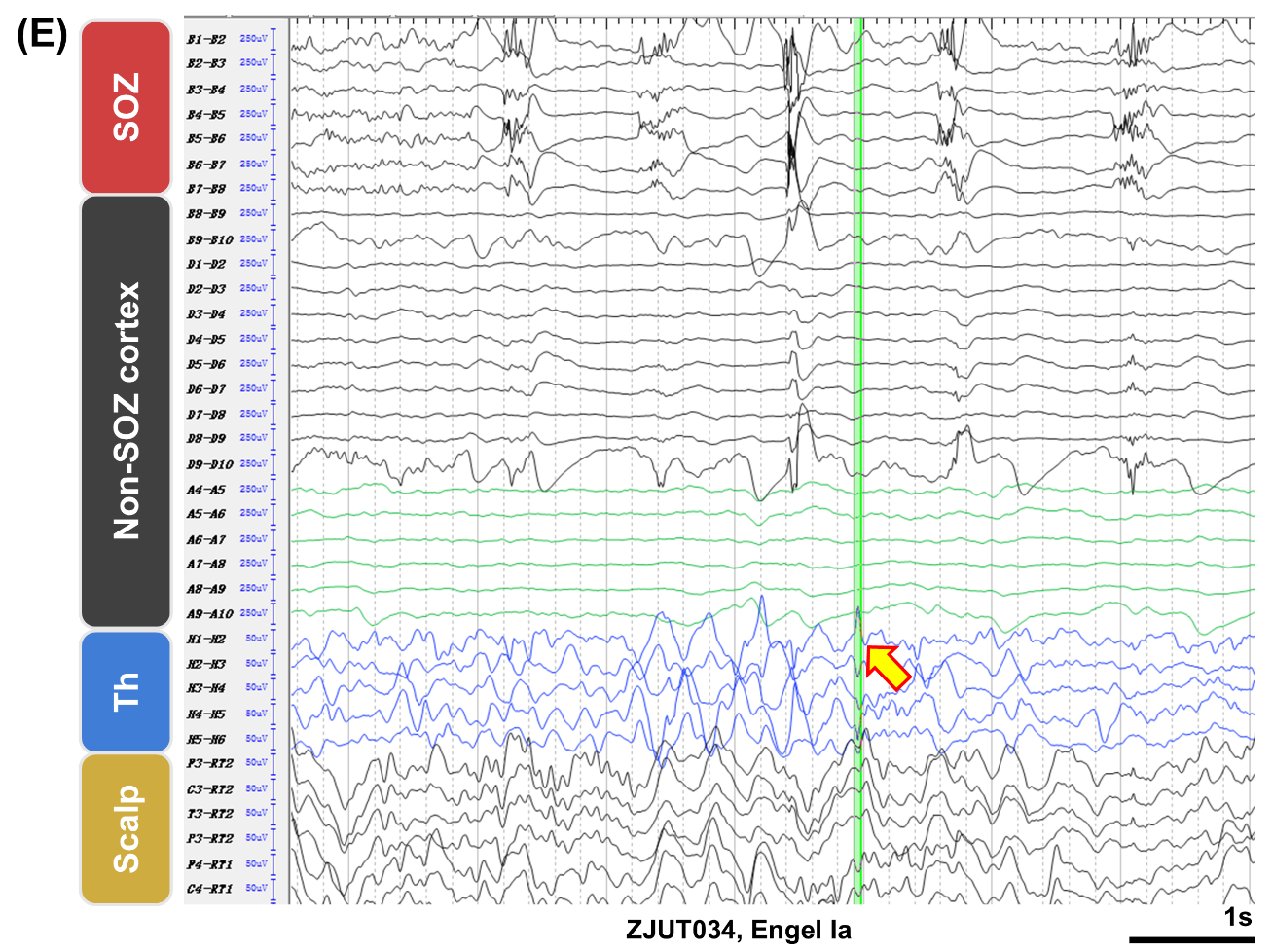

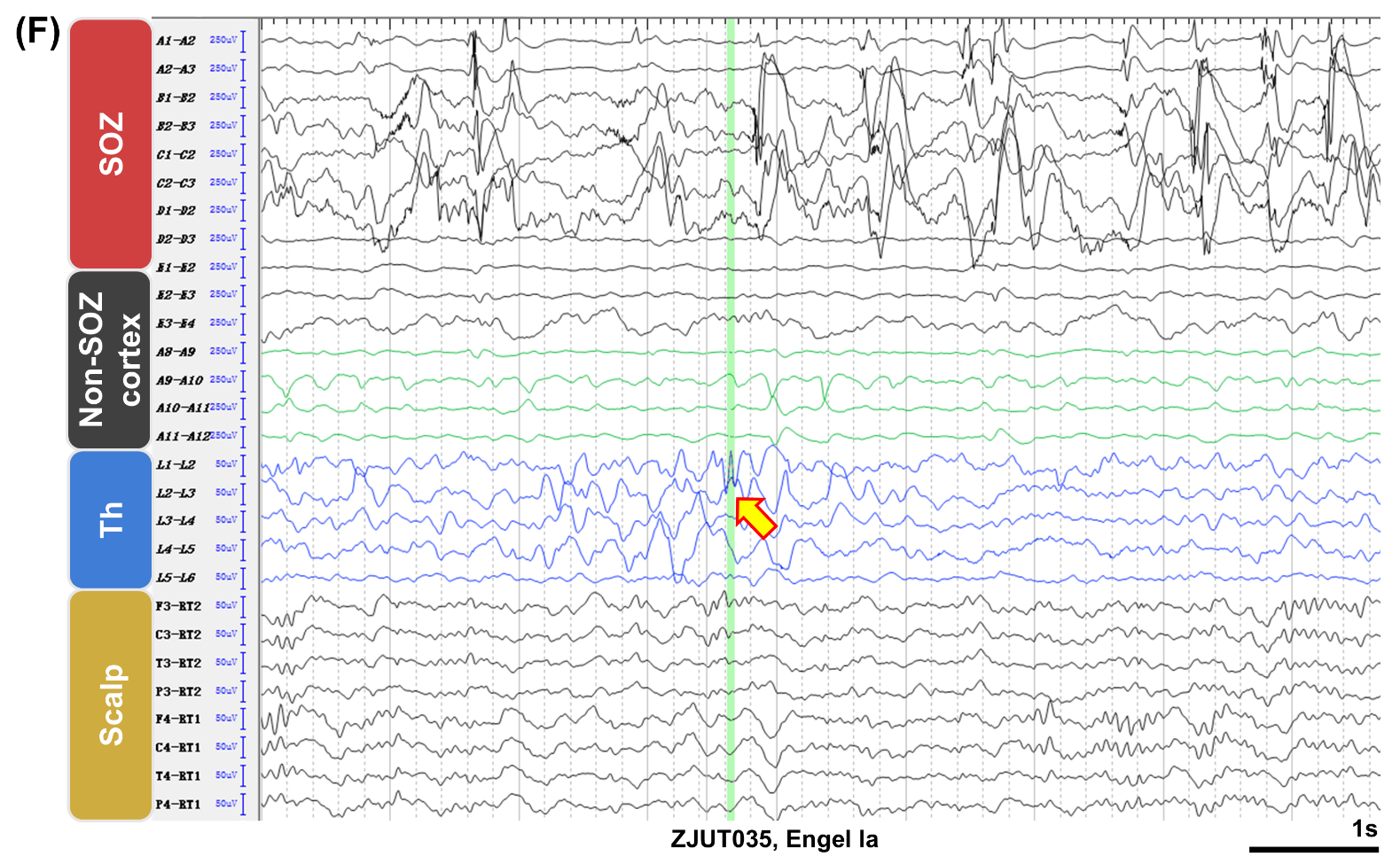

**Figure S13**. Examples of isolated spikes in the thalamus. Some events with the spike morphology were observed in the thalamus but they did not co-occur with cortical spikes. They could be either physiological activity or epileptiform discharges propagated from SEEG unsampled areas. Intracranial signals were filtered between 0.2 and 250 Hz, and scalp signals were filtered between 0.2 and 70 Hz. Patient numbers and surgical outcomes are presented at the bottom. The "RT" channel in the scalp is located in the mastoid. (A) The SOZ of this patient was located in the mesial temporal lobe, and the pathology was **hippocampal sclerosis**. (B) The SOZ of this patient was located in the anterior cingulate cortex, and the pathology was gliosis. (C) The SOZ of this patient was located in the anterior insula and operculum, and the pathology was **FCD IIa**. (D) The SOZ of this patient was located in the mesial frontal lobe, and the pathology was **FCD IIb**. (E) The SOZ of this patient was located in the posterior insula and operculum, and the pathology was **FCD II**. (F) The SOZ of this patient was located in the mesial temporal lobe, and the pathology was gliosis. SOZ, seizure onset zone; Th, thalamus; FCD, focal cortical dysplasia.

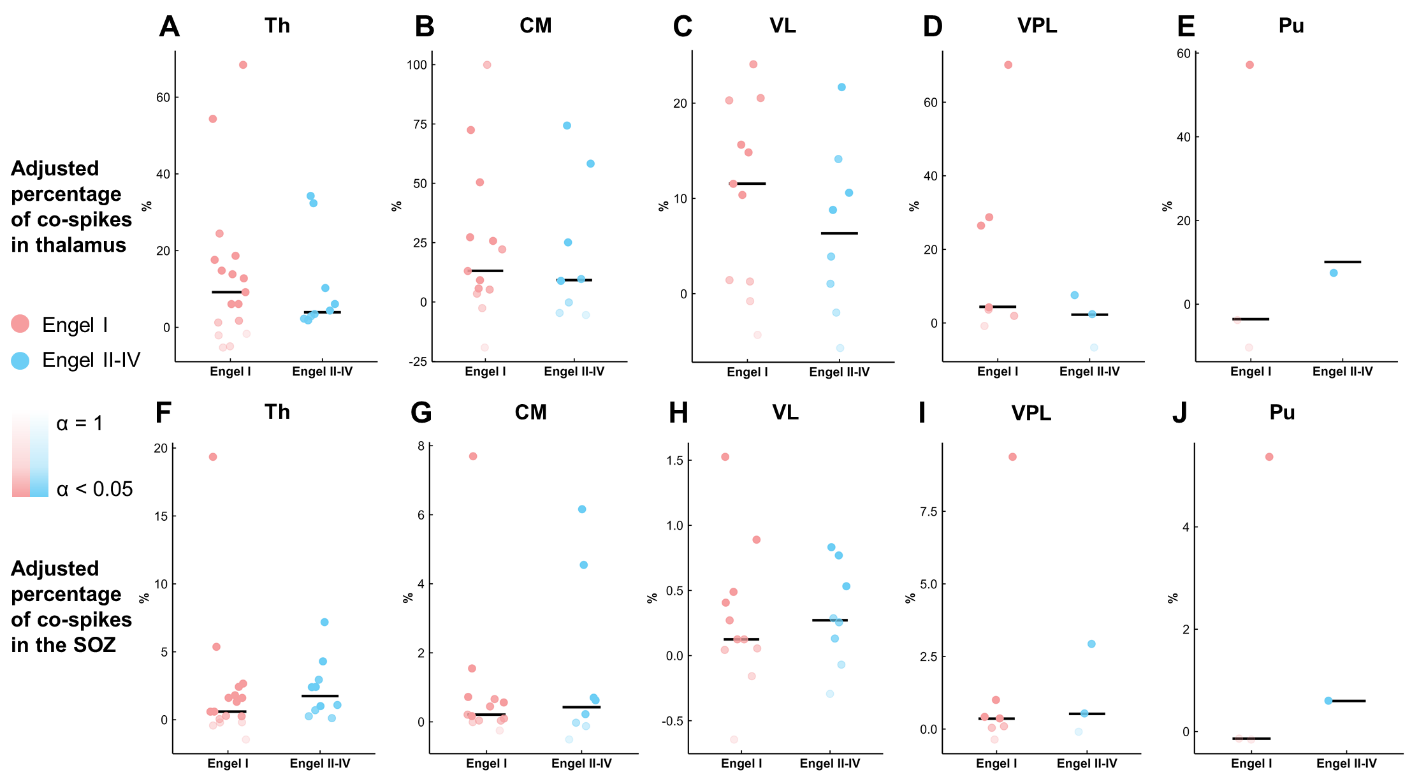

**Figure S14.** The proportions of co-spikes between the SOZ and thalamus in patients with focal SOZs and their relationships with surgical outcomes **when using the raw output from the automated spike detector**. (A)-(E) The percentage of co-spikes obtained when the number of thalamic spikes was used as the denominator. Most of thalamic spikes do not co-occur with spikes in the SOZ. (F)-(J) The percentage of co-spikes obtained when the number of SOZ spikes was used as the denominator. SOZ spikes uncommonly propagate to the thalamus. There was no difference between favorable and unfavorable surgical outcome group. Each dot represents one subject. The transparency of the color indicates the alpha value in the surrogate data analysis. Lower transparency means that the number of co-spikes is higher than the chance level. SOZ, seizure onset zone; Th, thalamus as a whole; CM, centromedian nucleus; Pu, pulvinar nuclei; VL, ventral lateral nucleus; VPL, ventral posterolateral nucleus.

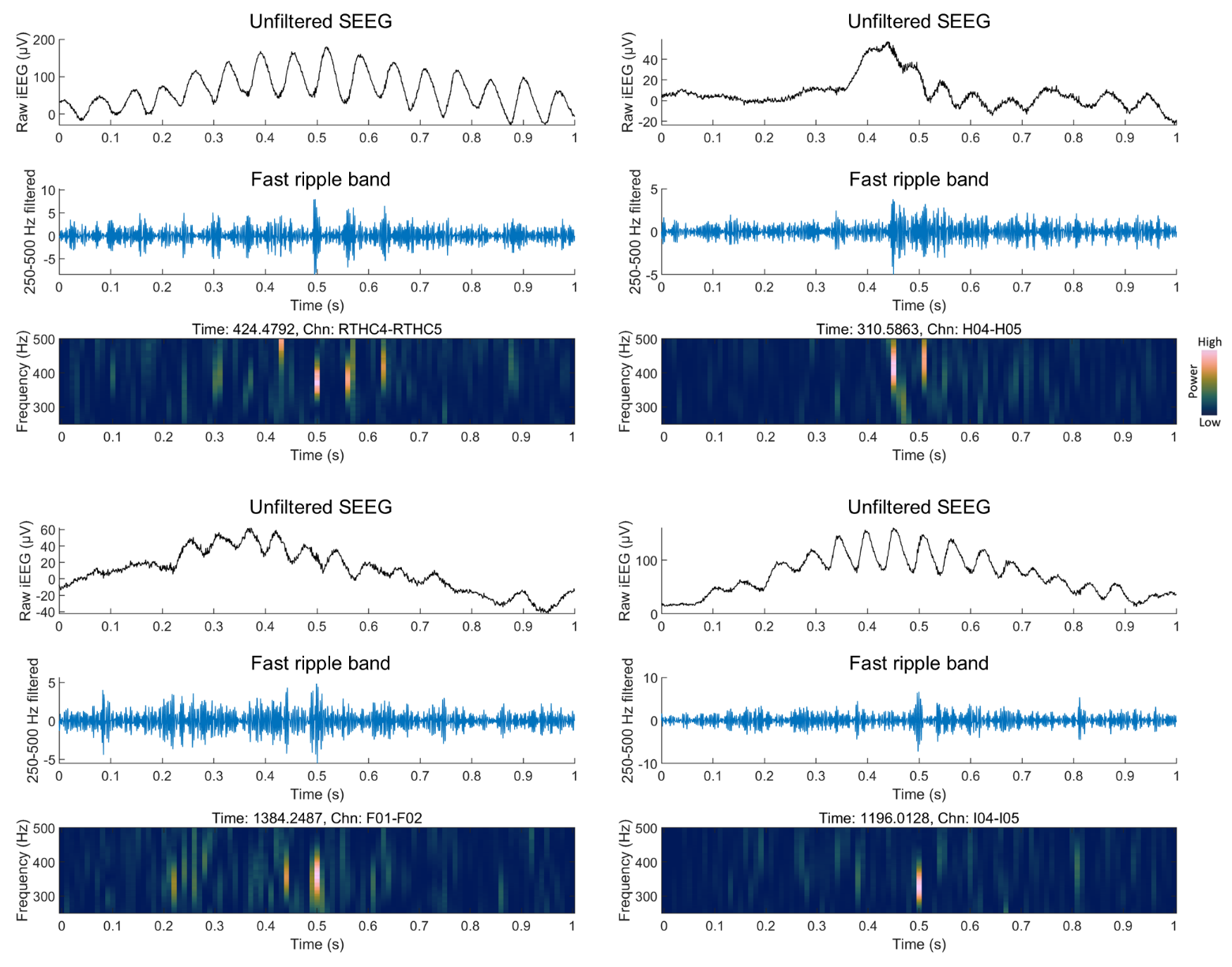

**Figure S15.** Examples of fast ripples superimposed on sleep spindles in the human thalamus. Signals and time-frequency spectra are showed in the figure. It can also be observed that **fast ripple band activity (250-500 Hz) is likely to have phase amplitude coupling with sleep spindles**. SEEG, stereo-electroencephalography; Chn, channel number.
